# Evolution of CRISPR-associated Endonucleases as Inferred from Resurrected Proteins

**DOI:** 10.1101/2022.03.30.485982

**Authors:** Borja Alonso-Lerma, Ylenia Jabalera, Matias Morin, Almudena Fernandez, Sara Samperio, Ane Quesada, Antonio Reifs, Sergio Fernández-Peñalver, Yolanda Benitez, Lucia Soletto, Jose A Gavira, Adrian Diaz, Wim Vranken, Benjamin P. Kleinstiver, Avencia Sanchez-Mejias, Marc Güell, Francisco JM Mojica, Miguel A Moreno-Pelayo, Lluis Montoliu, Raul Perez-Jimenez

## Abstract

Clustered regularly interspaced short palindromic repeats (CRISPR)-associated Cas9 protein is an effector that plays a major role in a prokaryotic adaptive immune system, by which invading DNA can be targeted and cut for inactivation. The Cas9 endonuclease is directed to target sites by a guide RNA (gRNA) where Cas9 can recognize specific sequences (PAMs) in foreign DNA, which then serve as an anchoring point for cleavage of the adjacent RNA-matching DNA region. Although the CRISPR-Cas9 system has been widely studied and repurposed for diverse applications (notably, genome editing), its origin and evolution remain to be elucidated. Here, we investigate the evolution of Cas9 from resurrected ancient nucleases (anCas) in extinct firmicutes species as old as 2600 myr to the current day. Surprisingly, we demonstrate that these ancient forms were much more flexible in their PAM and gRNA scaffold requirements compared to modern day Cas9 enzymes. In addition, anCas portrays a gradual paleoenzymatic adaptation from nickase to double-strand break activity, suggesting a mechanism by which ancient CRISPR systems could propagate when harboring Cas enzymes with minimal PAMs. The oldest anCas also exhibit high levels of activity with ssDNA and ssRNA targets, resembling Cas nucleases in related system types. Finally, we illustrate editing activity of the anCas enzymes in human cells. The prediction and characterization of anCas proteins uncovers an unexpected evolutionary trajectory leading to ancient enzymes with extraordinary properties.

## Introduction

The CRISPR-Cas molecular complexes of prokaryotes are very diverse in function^1^ and composition^2^, with currently in over thirty subtypes categorized in six types and two classes (types I, III and IV in class 1, types II, V and VI in class 2). Most of them serve as defense systems that prevent infection by viruses and other transmissible genetic elements. To defend against foreign nucleic acids, CRISPR loci harbour repeat-intervening DNA fragments (spacers) acquired during the so-called adaptation stage from invading nucleic acid fragments that have attacked the cell lineage in past generations^3–6^. In many CRISPR-Cas systems, only DNA fragments next to specific short sequences called protospacer adjacent motifs (PAMs)^7^ are incorporated as new spacers. Over time, adaptation has created a vast collection of invading DNA spacers that are continuously evolving^8^.

Transcripts from the CRISPR array are processed into small RNA molecules (crRNA), each containing a fragment of a single spacer and part of the repeats. The following steps of the CRISPR mechanism may show substantial variations depending on the system type^9^. In the type II CRISPR-Cas system, associated with the signature Cas9 effector protein (referred to as the CRISPR-Cas9 system), the crRNA molecules bound to an accessory trans-activating crRNA (tracrRNA), serve as cognate guides (gRNAs) for the Cas9 nuclease. After binding to a compatible PAM in the invading dsDNA, Cas9 uses its RuvC and HNH nuclease domains^6^ to catalyse a double-strand break (DSB) within PAM-adjacent sequences that match the spacer carried by the crRNA.

The accuracy of the targeting depends on the binding of Cas9 to the gRNA, and the Cas9-gRNA complex subsequently to the PAM and remainder of the target site^10, 11^. This recognition ability is also the basis for the utilization of CRISPR-Cas9 as a gene editor, often linking the crRNA and tracrRNA into a single-guide RNA (sgRNA)^6, 12, 13^. The incorporation of invading DNA in the CRISPR locus is an adaptive process that is time-dependent, i.e., over time bacteria will acquire new DNA that will be transferred to new generations. The ability of the organisms to process new DNA for self-protection must have arisen due to evolutionary pressure, which might be responsible for the recognition and cleavage properties of Cas nucleases. For instance, although the absence of the PAM in the spacer-proximal sequence of the type II CRISPR repeats protects the CRISPR locus from targeting, the acquisition of spacers matching the host genome has the potential to induce autoimmunity^14^ and diverse mechanisms have been identified that prevent or minimize the deleterious consequences of CRISPR-driven DSB in prokaryotes^15^. However, how nucleases have achieved such abilities, as well as the origin of the system, is a matter of debate^16^.

In this work, we resurrect ancestral forms of Cas9 to demonstrate how evolution has likely sculpted Cas endonucleases. Ancestral proteins have demonstrated outstanding properties depicting their evolutionary history, as well as environmental and organismal characteristics^17, 18^. Importantly, numerous ancestral enzymes have shown abilities useful for biotechnological applications^19, 20^, making Ancestral Sequence Reconstruction (ASR) technique a powerful enzyme design tool with multiple purposes that can go beyond the capabilities of other protein engineering techniques^21^. Here, we have resurrected five ancestors of Cas9 (anCas), from a 2600 million years ago (mya) old common ancestor of a set of modern firmicutes (FCA) tracing an evolutionary path to modern *Streptococcus pyogenes* Cas9 (SpCas9). Unlike SpCas9, the oldest anCas can be guided by diverse sgRNA scaffolds and exhibits remarkable PAM flexibility, indicating that Cas9 evolved from total PAMless to 5’-NGG-3’ specificity. The activity of the anCas proteins depicts a paleotrend for nuclease/nickase activity ratio and PAM recognition that confers special abilities for gene targeting and cleavage. Furthermore, we demonstrate robust genome editing activity with these anCas in human cells. Overall, the anCas not only shed light onto the evolution and appearance of the CRISPR-Cas9 system, but also show unprecedented abilities for PAM and sgRNA recognition, nickase and nuclease activity, functional promiscuity, immune response and editing of human cells that makes them a valuable addition to the growing list of Cas nucleases with novel properties^22^.

## Results

### Reconstruction of ancestral nucleases: anCas

To perform a reconstruction of ancestral nucleases, we searched the NCBI Protein database using the sequence of subtype II-A CRISPR-associated endonuclease Cas9 (formerly Csn1) from *S. pyogenes* as query (NCBI Reference Sequence WP_032464890.1 and UniProt Q99ZW2) with full annotation score. The NCBI database offers a BLAST search that allows an initial sequence homology search using the Blosum62 scoring matrix. We retrieved a total of fifty-nine Cas9 sequences from firmicutes and actinobacteria species (Sequences in Supplementary Information). We constructed a sequence alignment that was used to generate a gene phylogenetic chronogram using Bayesian inference and Markov Chain Monte Carlo (MCMC) (**Fig. 1a**). Several internal nodes were selected for laboratory resurrection of anCas, tracing the evolutionary path from the most recent Firmicutes Common Ancestor (FCA) of the set, that lived in the Neoarchean era around 2600 mya, to modern *S. pyogenes*. This path also contained anCas from a Bacilli Common Ancestor (BCA) that lived in the Neo-Proterozoic era around 1000 mya, and anCas from a Streptococci Common Ancestor (SCA) from the Triassic period around 200 mya. Two other ancestors of SpCas9 were reconstructed, Pyogenic Common Ancestor (PCA) and Pyogenes-Dysgalactiae Common Ancestor (PDCA) that according to our tree calibration lived around 137 and 37 mya, respectively (**Fig. 1a**). Using maximum likelihood, the most probable ancestral sequences for these nodes can be reconstructed and synthetically resurrected (Online Methods).

**Figure 1.**
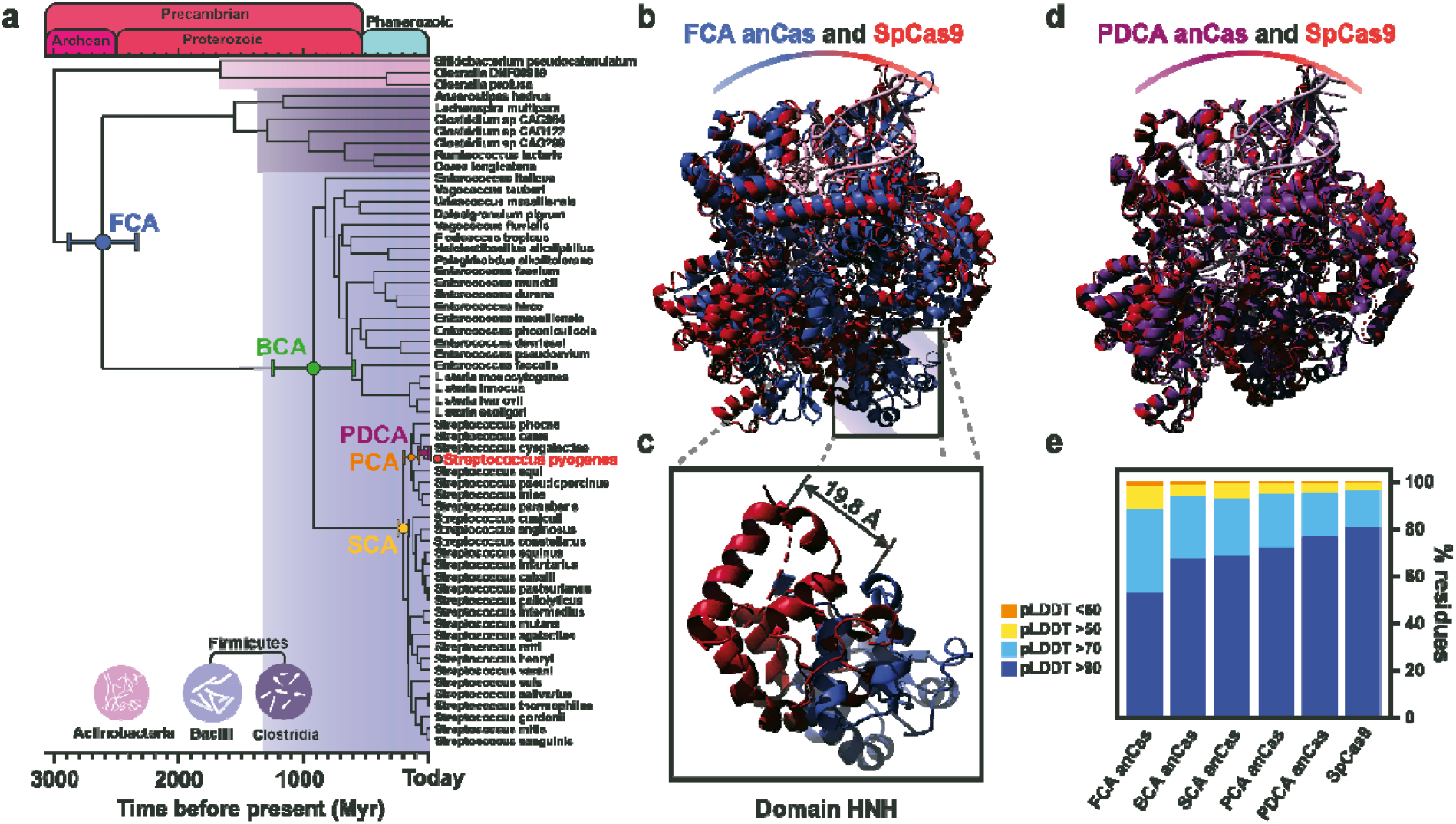
Phylogenetic and structural analysis of anCas endonucleases. **(a)** Phylogenetic chronogram of Cas9 endonucleases. Fifty-nine sequences were chosen from two phyla, Firmicutes and Actinobacteria, with two classes, Bacilli and Clostridia, belonging to Firmicutes. Identification codes of all sequence can be found in the Supplementary Information. Divergence times were estimated using Bayesian inference and information from the Time Tree of Life. Internal nodes from Firmicutes Common Ancestor (FCA), Bacilli Common Ancestor (BCA), Streptococci Common Ancestor (SCA), Pyogenic Common Ancestor (PCA) and Pyogenes-Dysgalactiae Common Ancestor (PDCA) were selected for testing. **(b)** Superposition of structural prediction of FCA Cas9 using AlphaFold2 (blue) with x-Ray structure of SpCas9 with guide RNA and target DNA (PDB:4oo8, red). **(c)** Isolated HNH domains of SpCas9 (red) and FCA anCas (blue) from whole structure coordinates. **(d)** Superposition of structural prediction of PDCA anCas with guide RNA and target DNA using AlphaFold2 (purple) with x-Ray structure of SpCas9 (PDB:4oo8, red). **(e)** pLDDT values for all anCas as estimated from AlphaFold2 prediction.

We compared the reconstructed sequences to that of SpCas9, yielding amino acid sequence identities in the range 53-96% (**Supplementary Fig. 1, 2**). The lowest identity corresponds to FCA anCas, which contains over 620 amino acid differences with respect to SpCas9. In the case of BCA and SCA anCas, the accumulated number of changes is 410 and 342 mutations, respectively, whereas for PCA and PDCA anCas only 226 and 54 mutations occur, respectively. Upon aligning all sequences, we observe that most important residues that have been reported to have a role in Cas9 function such as D10, S15, R66, 70, 74, 78, 1333 and 1335 and tandem residues such as PWN in 475-477 are all conserved across the anCas sequences when compared to SpCas9. However, some notable differences exist (**Supplementary Fig. 3**). Residue 1103, equivalent to W1126 in SpCas9, which seems to be associated to PAM recognition^23^, is mutated to L in FCA anCas. Similarly, amino acids in positions 1084 and 1306 in FCA anCas (corresponding to K1107 and K1334 in SpCas9) are mutated to D and M, respectively. Precisely, the tandem RKR in positions 1333-1335 located in the PAM-interacting (PI) domain has been shown to play a pivotal role in PAM recognition^10^. Finally, residue S1109, also associated to PAM interaction, is mutated to T in FCA and BCA anCas enzymes. These alterations suggest possible differences in PAM recognition abilities in FCA anCas and possibly in BCA anCas.

We used the anCas sequences to predict their structures using the newly developed AlphaFold2 structure prediction platform^24, 25^, which utilizes a neural network to infer structures with often near-experimental accuracy. Five structural models were predicted for each sequence and the model with the highest confidence was chosen for the structural analysis comparison (**Supplementary Fig. 4**). For all anCas, the estimated per-residue confidence score (Local Difference Distance Test), pLDDT^26^, is on average over 80 (**Fig. 1e, Supplementary Table 1**). We performed a structural alignment of each selected anCas model with the structure of SpCas9 bound to target DNA and gRNA (pdb:4oo8), (RMSD 2.52 Å) (**Fig. 1b, d**). This is expected given that most Cas9 structures available in databases are bound to target DNA through gRNA. Upon close examination of the structural alignment of FCA anCas and SpCas9, we observe a drastic structural difference in the position of the HNH domains, with a clear displacement of around 20 Å (red and blue domains in **Fig. 1c**). Considering that this domain is involved in DNA cleavage, an alteration in function would be expected. The rest of the anCas align well with SpCas9, with an increasing pLDDT as they approach modern days (**Fig. 1e, Supplementary Table 1**).

**Table 1.** Kinetics parameters of all anCas and SpCas9 determined from the single exponential decay curves in Figure 2c and 2e.

|  |  |  |  |  |  |  |  |  |  |  |  |  |
| --- | --- | --- | --- | --- | --- | --- | --- | --- | --- | --- | --- | --- |
| Kinetics parameters | SpCas9 |  | FCA anCas |  | BCA anCas |  | SCA anCas |  | PCA anCas |  | PDCA anCas |  |
|  | Total | DSB | Total | DSB | Total | DSB | Total | DSB | Total | DSB | Total | DSB |
| $k_{\text{cleave}}$ ($\text{min}^{-1}$) | 0.35 ± 0.01 | 0.194 ± 0.01 | 0.27 ± 0.03 | 0.0093 ± 0.0009 | 0.313 ± 0.003 | 0.019 ± 0.001 | 0.53 ± 0.02 | 0.16 ± 0.03 | 0.784 ± 0.003 | 0.53 ± 0.06 | 0.44 ± 0.01 | 0.24 ± 0.02 |
| Max fraction | 1 | 1 | 1 | 0.612 | 1 | 0.740 | 1 | 1 | 0.9994 | 0.89 | 1 | 1 |
| $R^2$ | 0.9994 | 0.9958 | 0.9910 | 0.9989 | 0.9999 | 0.9988 | 0.9998 | 0.9700 | 0.9999 | 0.9985 | 0.9997 | 0.9964 |

We also performed a comparative structural analysis of the different domains alone by calculating RMSD values (**Supplementary Table 2**). As expected, the highest RMSD values correspond to FCA anCas; however, direct domain comparison demonstrates considerable differences of over 2 Å in domains REC and RuvC III, which again further suggests that structural constraints might alter the function of FCA anCas. Surprisingly, when comparing the HNH domain alone between SpCas9 and FCA anCas, the RMSD is approximately 1 Å, demonstrating that the displacement observed in the complete structure (**Fig. 1c**) must be due to structural alterations to a global level in the FCA anCas structure. Finally, it is worth noting that RMDS values for PI domains seem to follow a decreasing trend, from the oldest to the newest anCas.

**Table 2.** Kinetics parameters of FCA and BCA anCas and SpCas9 determined from the single exponential decay curves in Figure 4e and 4f.

|  |  |  |  |  |  |  |
| --- | --- | --- | --- | --- | --- | --- |
| Kinetics parameters | SpCas9 |  | FCA anCas |  | BCA anCas |  |
|  | ssRNA | ssDNA | ssRNA | ssDNA | ssRNA | ssDNA |
| $k_{\text{cleave}}$ ($\text{min}^{-1}$) | 0.016 ± 0.009 | 0.002 ± 0.008 | 0.002 ± 0.01 | 0.045 ± 0.01 | 0.014 ± 0.006 | 0.045 ± 0.02 |
| Max fraction | 0.43 | 0.41 | 0.66 | 0.70 | 0.97 | 0.99 |
| $R^2$ | 0.9856 | 0.9902 | 0.9807 | 0.9824 | 0.9954 | 0.9529 |

### Activity of anCas nucleases: DSB and nickase activity

The anCas genes were synthesized and cloned into pBAD/gIII expression vector, carrying an arabinose inducible promoter and a *gIII* encoding signal that directs the anCas to the periplasmic space. All anCas were expressed at high levels in *Escherichia coli* BL21 cells.

We began to assess anCas activity test by assuming a simplistic scenario by which anCas would recognize a sgRNA from *S. pyogenes* as well as its canonical 5’-NGG-3’ PAM sequence (**Fig. 2a**). We designed a sgRNA containing a 20 nt-long spacer region targeted towards a DNA fragment upstream of a TGG PAM, all placed in a 4007 bp supercoiled plasmid. *In vitro* cleavage assays were carried out by incubating anCas or SpCas9 with target DNA together with the sgRNA at different digestion times. Although with clear differences in cleavage efficiency, all enzymes tested produce both relaxed and linear products, indicative of nickase and DSB activity, respectively. As expected, SpCas9 shows nicked product at short times and linear products at longer times (**Fig. 2b**). However, in the case of anCas the behavior changes from the oldest FCA anCas to more recent enzymes (**Fig. 2b**). FCA anCas mostly shows nickase activity and only at times over 60 min, the DSB activity becomes prominent. The other anCas show a progressive behavior with more intensive DSB activity in the younger anCas (**Fig. 2b**). We quantified both the nicked and linear fraction for each anCas and SpCas9 and plotted them versus incubation time in three forms, total cleavage (**Fig. 2c**), nicked fraction (**Fig. 2d**) and linear fraction (**Fig. 2e**), demonstrating the progressive decrease of nicked fraction and increase of linear fraction. SpCas9 has the higher proportion of linear products, the oldest FCA anCas has the highest proportion of nicked fraction (**Table 1**). We plotted the fraction of linear and nicked products at 30 minutes reaction versus geological time demonstrating an evolutionary trend from nickase to DSB activity (**Fig. 2f**).

**Figure 2.**
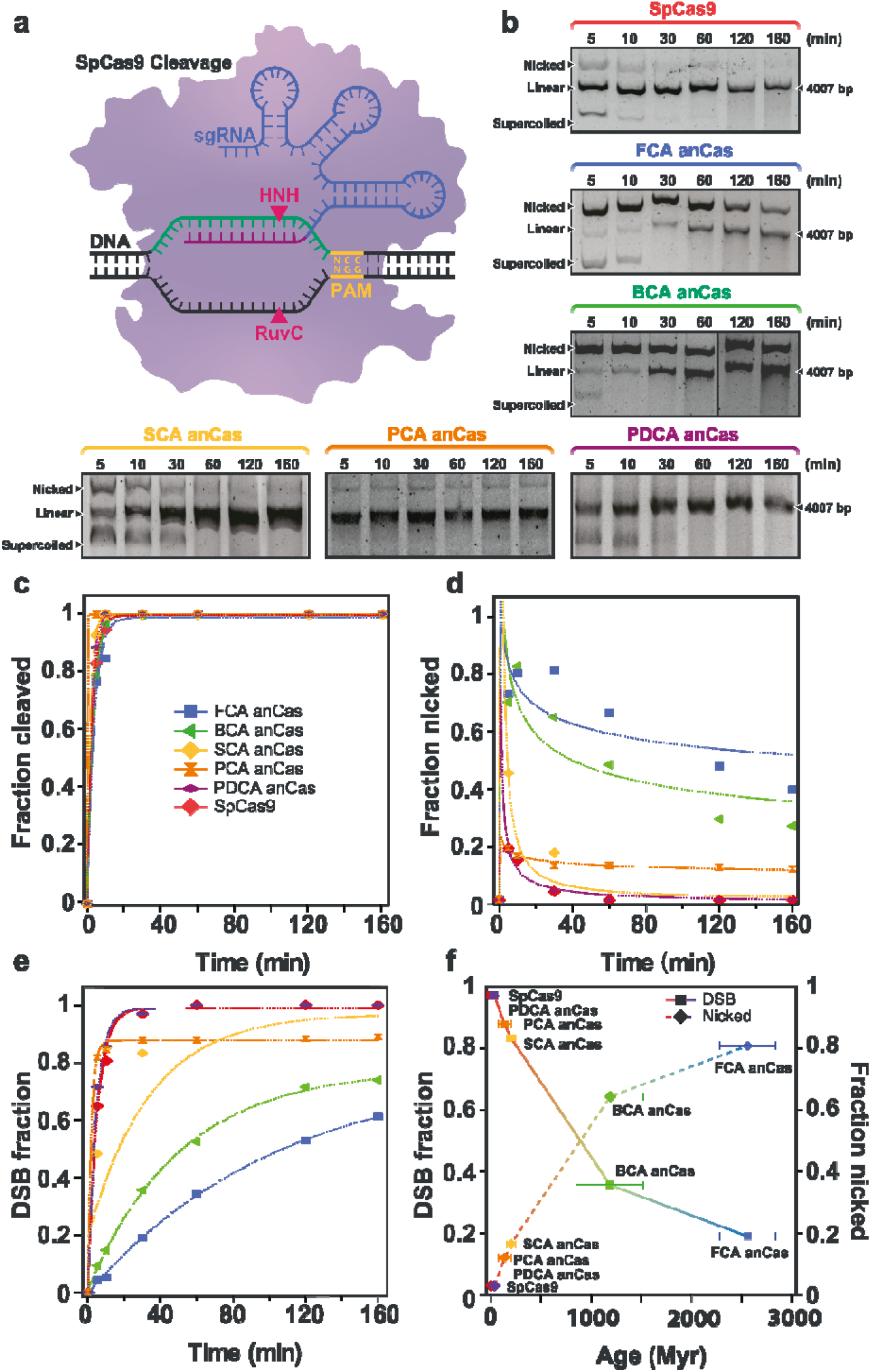
Activity of anCas endonucleases on a supercoiled DNA substrate. **(a)** Schematic representation of endonuclease activity on a supercoiled substrate. **(b)** *In vitro* cleavage assay for SpCas9 and all anCas on a 4007 bp substrate at different reaction times showing nicked and linear fractions. **(c)** Quantification of total cleavage at different reaction times and exponential fits (lines). **(d)** Quantification of nicked fraction for all anCas and SpCas9 at different times. **(e)** Quantification of DSB cleavage. Single-exponential fits were used to obtain *k*_cleave_ and maximum fraction cleaved (amplitude). Fitting parameters are summarized in Table 1. **(f)** DSB fraction (left axis) and nicked fraction (right axis) at 30 minutes reaction plotted against evolutionary time.

The structural differences observed in FCA anCas protein related to HNH domain displacement, together with the evolutionary trend from nickase to DSB activity, suggests the oldest anCas, FCA, could display an ancestral HNH domain with a reduced or suppressed activity. To examine this, we tested the *in vitro* activity of the H838A FCA anCas mutant (H840A with respect to SpCas9). The mutant was able to produce nicked and, surprisingly, linear products, showing a profile practically identical to that obtained with wild type FCA anCas (**Supplementary Fig. 5**). These results suggest that FCA anCas may contain an immature HNH domain, with the RuvC domain responsible for the nickase and DSB activity observed in FCA anCas, as has been previously shown in some type V effector nucleases that lack the HNH domain, such as Cpf1 (Cas12a), Cas14 (Cas12f) or CasF (Cas12j)^27–30^.

### Activity of anCas nucleases: PAM and sgRNA recognition

Compared to the more stringent gRNA scaffold and PAM preferences displayed by more evolutionary contemporary Cas enzymes, we were surprised that the anCas enzymes showed nuclease activity *in vitro* when using elements from SpCas9, i.e., gRNA and NGG PAM. We therefore investigated the ability of the anCas endonucleases to recognize different PAMs. To determine the preferred PAM sequence of each anCas, we cloned a DNA library containing a target sequence followed by seven random nucleotides (NNNNNNN) that corresponded to all possible PAMs (**Fig. 3a**). DNA libraries with random PAMs have been widely used in similar studies to characterize other Cas9 orthologs or mutated Cas9^31–33^. An sgRNA was designed using the scaffold of *S. pyogenes*. A 844 bp fragment containing both the target and PAM sequence was used as a substrate for anCas and SpCas9. We performed *in vitro* digestion using the purified Cas protein, and the transcribed sgRNA. All five anCas produced two fragments, one of 566 bp and a smaller one of 278 bp that would contain the PAM sequences recognized by the anCas (**Supplementary Fig. 6a)**. The small fragment was purified, sequenced by Next-generation Sequencing (NGS) and analysed to determine PAM sequence diversity from each anCas, enabling us to infer how evolution modified the PAM requirement of these enzymes. The five anCas enzymes and SpCas9 exhibited a range of PAM preferences displayed in the form of PAM wheels (Krona plot) (**Fig. 3b**). Surprisingly, FCA anCas exhibited no preference for any PAM sequence tested. For the other Cas proteins, we detected a preference for specific nucleotides in the target-proximal positions 2 and 3 (**Supplementary Fig. 6b and Supplementary Fig. 7**). Thus, in the case of BCA anCas, a slight preference for NGG was revealed, although additional PAM sequences were also detected (NNG). For the other, more recent anCas enzymes, the NGG bias was more prominent (**Supplementary Fig. 6b**).

**Figure 3.**
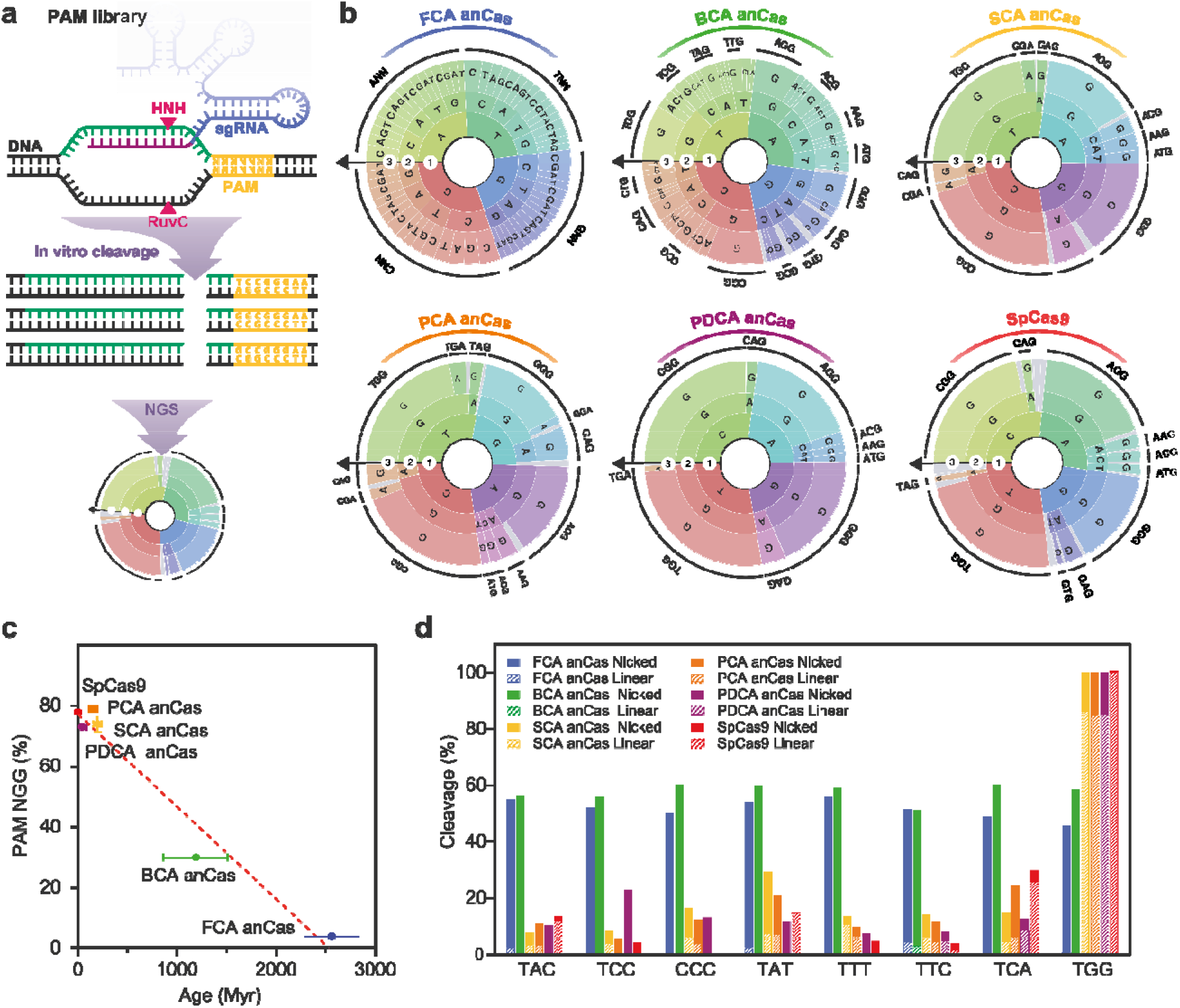
PAM determination of anCas. **(a)** Schematic representation of *in vitro* determination of PAM sequences using a library with random 7-nt PAM sequence using Next-generation Sequencing. Two fragments of 566 and 278 bp are generated after target cleavage. **(b)** PAM wheels (Krona plots) for all five anCas and SpCas9, used as control. **(c)** Percentage of reads containing an NGG PAM sequence 3-4 bp downstream the cleavage position plotted against evolutionary time. **(d)** *In vitro* cleavage assay (DSB and nicked products) using a variety of PAM sequences represented by TNN and CCC as control. Incubation time was 10 min.

After analysing all the sequences, we plotted the percentage of reads with an NGG PAM (**Supplementary Fig. 6c**) versus the geological time of each anCas, as estimated in the phylogenetic analysis. We observed a trend that reflects an NGG enrichment over time, demonstrating that NGG fidelity is an evolutionary trait that portrays a gradual progression from PAMless to NGG preference in more recent *Streptocococci* ancestors (**Fig. 3c**). This confirms the hypothesis of an evolving adaptive response for PAM recognition, which would be expected as the number of spacers acquired by the host cell increases over time. Eventually, a strong PAM recognition ability would be required to avoid self-cleavage of the CRISPR locus, especially, in a scenario in which the increment of DSB activity (deleterious in most prokaryotes) over nickase activity might further increase the evolutionary pressure on this ability. Although PAM-permissive Cas9 variants have been previously described^34–36^, to the best of our knowledge, FCA anCas is the first Cas9-related endonuclease that shows *per se* PAMless ability.

To further probe the PAMless ability of FCA anCas, we designed an *in vitro* PAM determination assay to test cleavage of a target DNAs adjacent to a total of seven selected PAM sequences of 3-nt length (TAC, TCC, TAT, TTT, TTC, TAC and TGG) within the general TNN PAM. A CCC PAM was also included in the set to verify possibilities other than an initial T nucleotide. anCas effectors together with the sgRNA were incubated with each of the target DNAs for 10 min and cleavage products were verified by agarose gel (**Supplementary Fig. 6d**). We observed both nicked and linear products, demonstrating the cleavage activity with all TNN PAM sequences. In the case of SpCas9, only TGG PAM demonstrated double-stranded break of the supercoiled DNA substrate. We quantified the percentage of nicked and linear products for each PAM, and observed that for the oldest anCas, FCA and BCA, the percentage of cleavage was similar for all PAM sequences tested, with mostly nicked products, as expected given the incubation time (**Fig. 3d**). In the case of the more contemporary anCas (PCA and DPCA) and SpCas9, the cleavage fraction reached high levels for TGG PAM, corroborating the NGG PAM preference. In the case of CCC control, the cleavage profile was similar to those obtained from non-NGG PAM sequences.

The promiscuity for PAM recognition exhibited by the oldest anCas made us wonder whether the anCas would also show promiscuity towards gRNA recognition as well. The reconstruction of an ancestral gRNA would have been interesting; however, the variability in sequence of crRNA repeats and tracrRNA from different species makes this challenging. To overcome this limitation and still evaluate the potential gRNA scaffold promiscuity of anCas, we decided to test modern sgRNAs from different species. We selected a total of five sgRNAs from *Streptococcus thermophilus, Enterococcus faecium, Clostridium perfingens, Staphylococcus aureus* and *Finegoldia magna*, covering several Firmicutes classes. These sgRNAs were selected following previous studies on sgRNA classification and function, in which sgRNAs were divided into seven clusters^37^. These distinct sgRNAs were contrasted against *S. pyogenes* guides containing spacers of two sizes, 18 and 20 nucleotides long, referred as 18 and 20 nt sgRNA, respectively.

SpCas9 and the five anCas were incubated during 10 min at 37 °C with a target plasmid DNA and TGG PAM recognition. We observed that, as expected, SpCas9 only linearized plasmid DNA when using its own sgRNA, although more efficiently when using the 20 nt spacer version, and sgRNAs from other species mostly resulted in nicked products leaving most supercoiled DNA substrate intact (**Fig. 4a**). On the contrary, FCA and BCA anCas were able to nick and linearize plasmid DNA with all sgRNAs, the *E. faecium* sgRNA showing better efficiency for FCA anCas, and the 18 nt sgRNA from *S. pyogenes* preferred for BCA anCas. We also tested the other anCas, observing that mostly FCA and BCA anCas had a marked promiscuity for sgRNA. All other anCas and SpCas9 seem to work best with 20 nt sgRNA from *S. pyogenes* (**Fig. 4b**).

**Figure 4.**
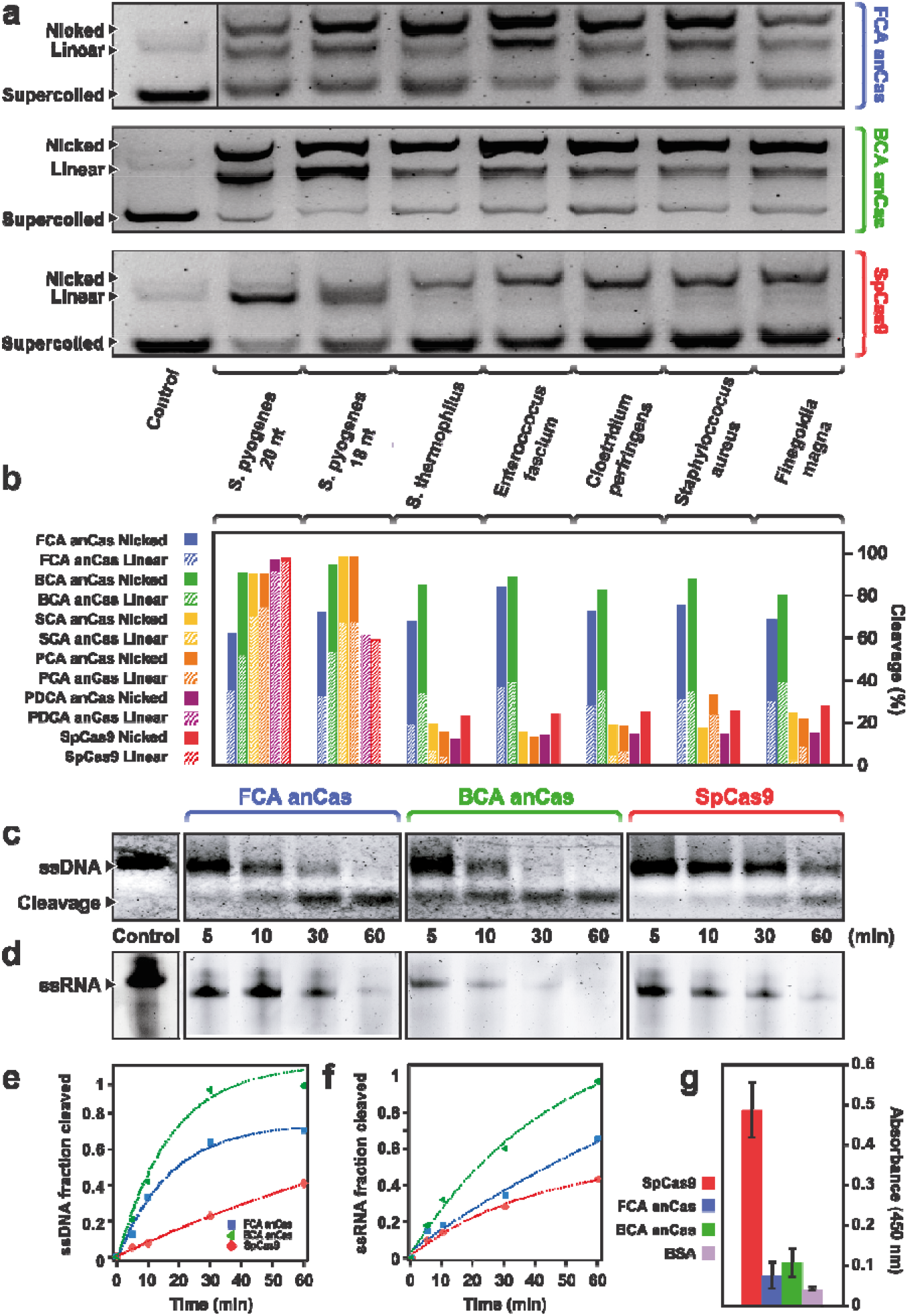
sgRNA test and nuclease activity of anCas on single-stranded substrates. **(a)** *In vitro* cleavage assay on a supercoiled DNA substrate of anCas and SpCas9 using sgRNAs from different species. FCA, BCA anCas and SpCas9 are shown. **(b)** Quantification of *in vitro* cleavage for all anCas and SpCas9 using the different sgRNAs. **(c)** *In vitro* cleavage assay on an 85 nt ssDNA fragment at different incubation times for FCA, BCA anCas and SpCas9. **(d)** *In vitro* cleavage assay on a 60 nt ssRNA at different incubation times for FCA, BCA anCas and SpCas9. **(e)** Quantification of fraction cleavage of ssDNA at different times and exponential fits for determination of kinetics parameters. In both (c) and (d), the control lane is the same for the three proteins. **(f)** Quantification of fraction cleavage of ssRNA at different times and exponential fits for determination of kinetics parameters. All kinetics parameters are summarized in Table 2. **(g)** Results from ELISA test of Anti-Cas9 rabbit antibody against SpCas9, FCA anCas, BCA anCas and BSA, used as control. Results are reported as average and S.D from three independent experiments.

Previous studies have indicated the contribution of the REC domain to sgRNA recognition specificity^38, 39^. As shown in Supplementary Table 2, this domain exhibits the highest RMSD differences, with a decreasing RMSD trend from the oldest to the newest anCas. These findings, along with the sgRNA promiscuity observed in oldest anCas, suggest that evolutionary pressure might have guided Cas nucleases to an enhanced guide specificity over time. In fact, this promiscuity has been already observed in type II-C Cas9, which has been suggested to be an ancient reminiscence of Cas9 nucleases^38^. In these nucleases, this promiscuity is also associated to PAM-independent ssDNA cleavage and weaker substrate DNA unwinding capabilities.

### Activity of anCas nucleases on single-stranded targets: ssDNA and ssRNA

As mentioned, the oldest anCas (FCA and BCA), show a remarkable nickase activity. This nickase activity may be related to ssDNA activity. ssDNA cutting activity has been suggested to be an ancestral trait present in smaller Cas9 such as subtype II-C Cas9^38^. This could also be reflected in the nickase activity of the ancestral forms from subtype II-A, such as anCas. Earlier forms of Cas9 with smaller catalytic domains might have been the origin of this ssDNA cutting activity that was still present in larger ancestral nucleases, which then gradually evolved towards DSB activity as part of a specialization process. To test this hypothesis, we examined the oldest anCas with an 85 nt ssDNA substrate containing target sequence complementary to the 20 nt spacer region of a SpCas9-sgRNA. As shown in **Fig. 4c, d**, FCA and, especially BCA anCas, show highest levels of ssDNA cleavage than those of SpCas9. Exponential fits to the data show much faster rates for FCA and BCA anCas, reaching almost full cleavage for BCA anCas (**Table 2**). We also tested activity on a 60 nt ssRNA target showing comparable results (**Fig. 4e, f**). Exponential fits of the cleavage activity reveal maximum rate and amplitude for BCA anCas, reaching again full cleavage (**Table 2**). These results demonstrate that both ancient Cas, but in particular, BCA anCas, behave as RNA-guided RNAses.

The activity of the oldest anCas on single-stranded substrates suggests that early Cas nucleases might have been active on those substrates, which seems to be an ancient trait, as mentioned before. These abilities may have additional important implication, given that the remarkable activity of anCas on ssDNA and ssRNA resembles that of Cas12a, Cas14 and Cas13a^27^, respectively, which suggest a connection among the activities of all class 1 effector nucleases. This functional promiscuity also puts anCas forward as a highly versatile endonuclease for genome editing applications.

We also investigated whether the oldest anCas endonucleases, which display more promiscuous features, may also have a different response towards an Anti-Cas9 antibody. We incubated BCA and FCA anCas with an Anti-Cas9 rabbit antibody and performed an ELISA test, which shows a diminished antibody binding (**Fig. 4g**). This would be expected given that host organisms carrying these nucleases have been long extinct and therefore have not been in contact with any living organisms. We believe that antibodies against Cas9 may have a weaker response towards ancient Cas forms. This lower antibody response might be of interest for potential applications in *in vivo* editing, where the immune response towards SpCas9 and other modern endonucleases represents a current limitation^40^.

### *In vivo* activity of anCas variants: determination of Indels

We then tested the genome editing activity of these ancestral nucleases in mammalian cells in culture, to answer the question whether these synthetic ancestral Cas can perform DNA cleavage; DSB, and trigger editing in cells under similar conditions as those associated with the standard SpCas9. To do so, the endogenous Tyrosinase gene (*TYR)* and the Melanocyte-specific transporter protein gene (*OCA2)*, whose mutations are associated to albinism^41^, were targeted in human HEK293T cells. The cells were co-transfected with plasmids containing the humanized versions of anCas or SpCas9, as well as the corresponding sgRNAs. We used standard sgRNA from *S. pyogenes* carrying a 20-nt spacer target. Seventy-two hours after co-transfection, cells were collected, and the genomic DNA extracted. The occurrence of indels (insertions/deletions) was first verified by the T7 endonuclease mismatch assay (**Fig. 5a**), where we could detect the expected DNA fragments. The size of the observed DNA fragments match those generated by DSB at the intended nucleotide, as well as the result of the subsequent cellular Non-Homologous End Joining (NHEJ) repairing pathway. T7-derived DNA fragments were obvious for the *OCA2* gene for all endonucleases except FCA anCas, whereas in the case of *TYR* locus the expected DNA bands were clearly visible for SpCas9 and PDCA anCas, and much weaker for SCA and BCA anCas (**Fig. 5a**). No bands were detected with the oldest anCas FCA in both loci.

**Figure 5.**
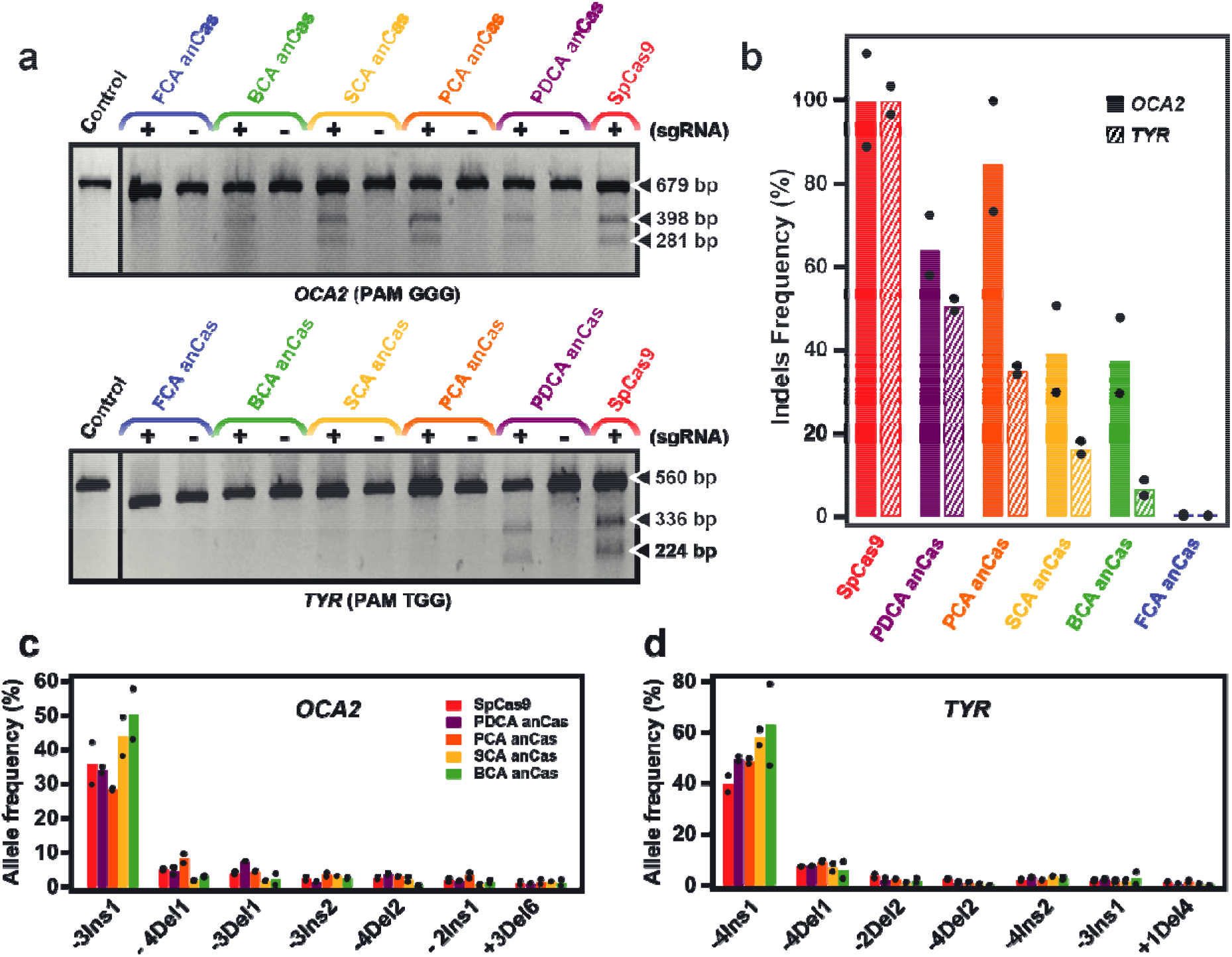
Activity of anCas endonucleases in HEK293T human cells. **(a)** T7 endonuclease mismatch assay for *OCA2* and *TYR* genes. Expected fragments for *OCA2* were 398 and 281 bp for a 679 bp amplified target, and 336 and 224 bp for *TYR* for a total 560 bp fragment. Experiments with (+) and without (-) sgRNA were run for each anCas. **(b)** Indels determination for *OCA2* and *TYR* by NGS of the experiments in (a) analyzed using Mosaic Finder. Indels frequency is normalized to SpCas9. **(c)** Allele frequency for *OCA2* and **(d)** *TYR*. Position of preferred cleavage was analyzed determining the preference, insertion (Ins) or deletion (Del) with position indicated upstream (-) or downstream (+) with respect to PAM. Length of the indel is indicated in the second figure (e.g. −3Ins1). The preferred allele for all anCas for *OCA2* is one insertion at three nucleotides upstream the PAM (−3Ins1). In the case of *TYR*, we find one insertion at the fourth nucleotide upstream the PAM as the preferred allele (−4Ins1). Bars represent the average value of two independent experiments indicated by the black dots.

We then evaluated and quantified indels by Next-generation Sequencing (NGS) using advanced analysis with Mosaic Finder software^42^, which allows high resolution reading of short DNA sequences after NHEJ repair. As shown in **Fig. 5b**, anCas endonucleases perform robust gene editing in human genomic DNA, except for FCA anCas. This is expected given the unique features of FCA anCas, which presumably does not use the HNH domain for cleavage and seems to function more optimally on single-stranded substrates, akin to other types of Cas nucleases. In addition, FCA anCas has shown a prominent nickase activity, which makes it a potential candidate for double-nicking editing. For gene *TYR*, a trend in indels proportion is observed (**Fig. 5b**). In the case of gene *OCA2* the proportion of indels reaches levels like those of SpCas9, e.g., PCA anCas, but even for oldest SCA and BCA anCas the levels are quite high, in the range of 40% (**Fig. 5b**). Comparable results were obtained analysing the NGS output files with the publicly available CrispRVariants R-software package^43^. Furthermore, the analysis of NGS reads also allows determination of preferred cleavage locations, as well as the appearance of the most frequent genome-edited alleles, whose pattern is maintained across all anCas that cleave these DNA sequences (**Fig. 5c, d and Supplementary Fig. 8**). This suggests that the outcome of the genome editing experiments, irrespective of the Cas used (anCas or SpCas9), is strongly determined by the surrounding genomic DNA sequence, as has previously been reported for SpCas9^44^.

Finally, we also tested site-specific cleavage using a Traffic Light Reporter (TLR) based on RFP reconstitution (see methods). This method allows monitoring of DNA repair in HEK293T cells based on fluorescence activated cell sorting (FACS). Using conditions optimized for SpCas9, the results are in line with those determined by NGS (**Supplementary Fig. 9**), demonstrating their robustness.

Overall, we demonstrate that anCas nucleases can potentially be used to edit cells; however, it is important that the conditions tested here are those favorable to SpCas9. We expect that searching other conditions involving, for instance, different sgRNA scaffolds, targets with various sequence contexts, nuclear location signal (NLS) configurations on the Cas protein, timing, etc, could lead to improvements allowing for anCas mediated gene editing in conditions beyond the current SpCas9 capabilities.

## Discussion

In the present study, we traced the evolutionary history of Cas9 endonucleases going back to the origin of Firmicutes species, where Cas9 is widely present. The ancestral variants, named anCas, display unique features not observed in their modern descendants. The evolutionary trends observed for the nuclease/nickase activity, NGG PAM recognition, gRNA recognition, and single-chain cleavage, highlight the adaptive character of the CRISPR-Cas systems, as well as offering features compatible with ancient functions. Characteristics such as PAM requirement or DNA unwinding are key to secure proper recognition of foreign versus self-DNA, as well as R-loop formation for cleavage. Ancient endonucleases likely resulted from enzymes that worked with ssDNA or RNA^38^, and did that not require a potent dsDNA unwinding ability, perhaps only needing additional proteins or co-factors to exert DSB activity^38^. The observation that the oldest anCas work better than SpCas9 with single-chain substrates demonstrates that they have good cleavage abilities but are perhaps less efficient at unwinding. Moreover, ancient Cas nickases that were not exposed to frequent viral invasions, and therefore did not have an extensive collection of spacers, likely did not need a strong PAM recognition; this is demonstrated by the oldest anCas.

The cleavage abilities of FCA anCas resemble those of other Cas enzymes such as Cas12a, Cas14 or CasF, in which cleavage results only from the action of the RuvC domain. These similarities suggest a common origin for all Cas enzymes, making them homologs^16^. This was already suggested for Class 2 CRISPR-Cas systems, which diverged to develop distinctive abilities after a specialization process. Features such as efficient PAM recognition and cleavage efficiency of dsDNA involving the HNH domain were already present in ancient *Streptococci* forms such as SCA, PCA and PDCA anCas. This suggests that these abilities where an important evolutionary trait that led to the divergence of modern *Streptococcus* spp. Overall, we can say that the most prominent feature of the oldest endonucleases is their chemical promiscuity for PAM, gRNA and substrate, which has been suggested as a common feature in early enzymes. Specificity is a chemical feature that in any enzyme is sculpted over millions of years of evolution driven by a continuous speciation and specialization process. This is exactly what we observed in younger anCas, where specificity and efficiency in dsDNA cleavage increases compared to the oldest anCas.

Ancestral Sequence Reconstruction (ASR) has demonstrated its validity to test evolutionary theories and its capacity to redesign enzymes with exceptional abilities based on their origins^18, 19, 45–49^. The excellent and abundant work on ancestral proteins and genes carried out in the past years has clearly placed this method as an exquisite biomolecular design technique that offers several advantages over existing techniques^20, 38, 50, 51^. Amongst the properties reported on ancestral proteins we can find increments in stability, functional promiscuity, pH tolerance, altered immunogenicity, structural alterations, and quite often, some or all these properties at once. Therefore, ASR should not be seeing as an exotic method for studying molecular evolution, but also as a method for exploring the sequence space of proteins by adding a time component that greatly expands the applicability of any protein. No other technique can currently yield a protein with hundreds of mutations with respect to existing ones, and still be fully folded and functional. These novel sequences, which no longer exist in nature, can thus be considered synthetic redesigned genes with known and unknown properties that, in most cases, can be explained under the light of evolution. Nevertheless, ancestral enzymes may also be the initial point for further improvement and modifications of proteins using rational design or directed evolution^50, 52^. Thus, ASR is a timely addition to the list of valid methods for protein design, providing perhaps the best possible approach for (re-)designing proteins and enzymes.

In the case of CRISPR-Cas, efforts have been made towards using evolutionary information to redesign Cas9 variants with the purpose of expanding their PAM range recognition, efficiency, and accuracy^34, 35^. Similarly, directed evolution approaches have been used to increase Cas9 specificity^53^. These new Cas9 have expanded the list of Cas9 that have been engineered for similar purposes^54, 55^. Still, the obtained variants are closely related orthologs whose sequences show only a few mutations, with most of them still showing limitations inherent to SpCas9. To circumvent CRISPR-Cas9 limitations, researchers have embarked on searches to identify new variants of Cas nucleases^22, 37^. This has resulted in an expanded catalogue of Cas nucleases that display a large diversity of molecular features. These Cas nuclease sequences provide the basis for the present study and allowed us to build a phylogenetic chronogram. Thus, discovering new Cas nucleases will provide new opportunities to expand the search in primordial time for their ancestors, further increasing the list of orthologs. Our study contributes the five anCas tested in this work, but also the sequences of the untested anCas nodes in the tree, which will likely work with known and, perhaps, new features.

The anCas tested in this study show a variety of features that make them potentially useful in genome editing applications. Altogether, the anCas variants have shown PAMless activity, gRNA promiscuity, nickase activity, marked activity on single-chain substrates, high cleavage efficiency and diminished immune response to an AntiCas9 antibody. All these features are desirable in emerging gene editing tools. In addition, the cleavage test in human cells demonstrates that most anCas can perform DSB followed by NHEJ in living cells, highlighting the validity of the ancestral reconstruction approach for the design of novel endonucleases, beyond the evolutionary implications. In addition, the prominent nickase activity of the oldest Cas such as FCA, makes it a potential candidate for double-nicking editing. Thus, this work represents a proof of concept that opens new avenues towards expanding the catalog of genome editing tools. Exploring these editing abilities including off-targets analysis, double-nicking cleavage, prime editing, complex DNA regions or other specific tests for biotechnology applications are clear directions for future work.

Finally, we believe anCas nucleases may be basic elements in which we can follow a modular Lego-like approach to fuse structural and molecular elements to redesign synthetic nucleases, or even combine them with previously developed Cas variants. Combining ASR and rational design would expand even further the sequence space, making these synthetic enzymes the basis of a collection of nucleases created *à la carte*. We also believe that any nuclease, as well as any enzyme, offers the potential to be reconstructed, offering countless possibilities not only for gene editing, but also for biotechnology in general.

## Supporting information

Evolution of CRISPR_SI

## Methods

### Ancestral sequence reconstruction of Cas9

Fifty-nine Cas9 sequences were downloaded from the NCBI database (Supplementary Information). Sequences belong to three bacterial phyla: Firmicutes, Clostridia and Actinobacteria. Alignment of the sequences was performed using MUSCLE software on the MEGA platform and manually edited. We inferred the best evolutionary model using MEGA, resulting in the Le and Gascuel (LG)^56^ with gamma distribution model (8 categories), Yule model for speciation and length chain of 20 million generations, sampling every 1000 generations. Phylogeny was carried out using BEAST v1.10.4 package software (https://beast.community/) including the BEAGLE library for parallel processing and based on Bayesian inference using Markov chain Monte Carlo (MCMC). We set *Streptococci* species as monophyletic group. The divergence times were estimated by uncorrelated log-normal clock model (UCLN), using molecular information from the Time Tree of Life (TTOL, http://www.timetree.org/) with default birth and death rates. Calculations were run in a multicore server. From the generated trees we discarded the initial 25% as burn-in using the LogCombiner utility from BEAST. We verified the MCMC log file using TRACER and ensure all parameters showed effective sample size (ESS)>100. Posterior probabilities of all nodes were above 0.65 and most of them were near 1. Figtree v1.4.3 was used to visualize and edit the phylogenetic tree. Finally, ancestral sequence reconstruction was performed by maximum likelihood using PAML v4.9 (http://abacus.gene.ucl.ac.uk/software/paml.html) with a gamma distribution for variable replacement rates across sites. Posterior probabilities were calculated for all amino acids and the residue with the highest posterior probability was chosen for each site. We selected from the tree to reconstruct different ancestors with different age named as anCas. These ancestors included an evolutionary line connecting an early ancestor of Firmicutes with modern *Streptococcus pyogenes*.

### Structure prediction of anCas endonucleases using AlphaFold2

The AlphaFold2 models^24, 26^ were calculated on the Flemish supercomputer centre (VSC) cluster at the University of Gent. The full AlphaFold2 pipeline was employed, with the main inference step executed five times, besides standard MSA search, template search, and constrained relaxation. pLDDT values were in turn mapped to the B-factor column of the corresponding coordinate model using a custom Python script. Visualization of the resulting models, with colouring according to pLDDT values (from 0 to 100), was performed with PyMol (The PyMOL Molecular Graphics System, Version 2.0 Schrödinger, LLC.). The spectrum bar was generated using the spectrumBar.py script (<u>PyMOLWiki</u>). Protein superposition was done with the SUPERPOSE software^57^ of the CCP4 program suite^58^.

### Protein production and purification

anCas genes were synthesized, and codon optimized for *E. coli* cell expression. All anCas genes were cloned into pBAD/His expression vector (ThermoFisher Scientific) and transformed in *E. coli* BL21 (DE3) (Life Technologies) for protein expression. SpCas9 gene was purchased from addgene (Plasmid #62934). Cells were incubated in LB medium at 37□°C until OD600 reached 0.6, L-arabinose was added to 0.1% to anCas. IPTG was added to SpCas9 to 1□mM concentration for protein induction overnight at 20 °C. Cells were pelleted by centrifugation at 4000□rpm. Pellets were resuspended in extraction buffer (20 mM Tris-HCl pH 7.9, 200 mM NaCl, 25 mM Imidazole, 0.5 mM TCEP). 100 mg/mL of lysozyme (Thermo Scientific) was added to the pellet and incubated for 15 min. Then pellet was sonicated for 3 cycles for 10 min at 30% amplitude. Cell debris was separated by ultracentrifugation at 33,000 x g for 1□h. For purification, the supernatants were mixed with His GraviTrap affinity column (GE Healthcare) and eluted in elution buffer (20 mM Tris-HCl pH 7.9, 200 mM NaCl, 300 mM Imidazole, 0.5 mM TCEP). Fractions were collected and loaded to a HiTrap Heparin HP column (GE Healthcare) and eluted using a linear gradient of 0.2-2 M NaCl. Proteins were further purified by size exclusion chromatography using a Superdex 200 HR column (GE Healthcare) and eluted in 20 mM Tris-HCl pH 7.9, 200 mM NaCl. For protein purification verification sodium dodecyl sulfate–polyacrylamide gel electrophoresis (SDS–PAGE) was used with 8% gels. The protein concentration was calculated by measuring the absorbance at 280□nm in Nanodrop 2000C.

### sgRNA synthesis

sgRNA with the complementary sequence (Supplementary Information) to the target was synthesized and cloned into pUC18 vector. sgRNA sequence was amplified by PCR using Phusion® Hot Start Flex DNA Polymerase (NEB). PCR product was purified using mi-PCR Purification Kit (Metabion). sgRNA was synthesized using HiScribe T7 High Yield RNA Synthesis Kit (NEB). The PCR fragment present the T7 promoter at 5’ end and the sequence from sgRNA of *S. pyogenes* at 3’ ends. The reaction was incubated overnight, and gRNA was purified following the protocol of the kit Monarch® RNA Purification Columns. sgRNA integrity was analyzed by electrophoresis on 2% agarose gel on TBE buffer.

### *In vitro* cleavage assay

*In vitro* cleavage assay was performed with purified anCas and SpCas9 endonucleases. In all assays, 30 nM enzyme was incubated for 15 min with 30 nM gRNA at 1:1 ratio in the cleavage buffer (100 mM NaCl, 50 mM Tris-HCl, 10 mM MgCl_2_, 100 µg/BSA, pH 7.9) at 37 °C. Then, 3 nM target DNA (plasmid DNA containing TGG PAM) was added and incubated at different times depending on the experiment. Reaction was stopped by adding 6X loading dye (NEB) with EDTA and run 2% agarose gel. Gels were dyed with SYBR gold (ThermoFisher Scientific) and imaged with ChemiDoc XRS + System (Bio-Rad). Cleavage was quantified by ImageJ.

### PAM library construction

DNA library containing seven random nucleotides was designed and cloned into pUC18 plasmid by Genscript. This random library was transformed in XL1blue *E. coli* and amplified several times to achieve the maximal variability in the PAM sequences. A PCR fragment of 844 bp was amplified using the primers (Supplementary Material) from the DNA library containing the 7 random nucleotides.

### PAM determination

PAM determination assay was performed incubating in cleavage buffer: 3 nM of PCR fragment from the DNA library with 30 nM of each anCas and SpCas9 and 30 nM of gRNA targeting the 20 nucleotides upstream the 7 random nucleotides. The reaction was incubated for 1 hour at 37 °C and stopped by adding 6X loading dye (NEB) with EDTA and run 2% agarose gel. Gels were dyed with SYBR gold (ThermoFisher Scientific) and imaged with ChemiDoc XRS + System (Bio-Rad). The small fragment of 278 bp was purified from the agarose gel with GeneJet Gel Extraction kit (ThermoFisher Scientific). The fragment was sequenced by Illumina Sequencing and the obtained reads were mapped to the reference sequence using Geneious Prime (2020 version). The reads that aligned to the reference with zero mismatches were selected and the frequency was calculated for each PAM. From the frequency of each PAM we generate PAM wheels following previously published methods^59^.

For *in vitro* cleavage of specific PAM we cloned into Zero-Blunt TOPO plasmid different DNA fragment carrying each PAM (Supplementary Material). The cleavage assay was performed in cleavage buffer (100 mM NaCl, 50 mM Tris-HCl, 10 mM MgCl_2_, 100 µg/BSA, pH 7.9) at 37 °C. 3 nM of anCas and SpCas9 were incubated for 15 min with 3 nM gRNA at 1:1 ratio in cleavage buffer and 3 nM DNA plasmid was added. After 10 min, reaction was stopped by adding 6X loading dye (NEB) with EDTA and run 2% agarose gel. Similarly, gels were dyed with SYBR gold (ThermoFisher Scientific) and imaged with ChemiDoc XRS + System (Bio-Rad). Cleavage was quantified by ImageJ.

### *In vitro* cleavage assay for gRNA promiscuity

For *in vitro* cleavage for gRNA promiscuity, DNA plasmid carrying TGG PAM was used. The cleavage assay was performed in cleavage buffer (100 mM NaCl, 50 mM Tris-HCl, 10 mM MgCl_2_, 100 µg/BSA, pH 7.9) at 37 °C. 3 nM of anCas and SpCas9 were incubated for 15 min with 3 nM sgRNA of each species at 1:1 ratio in cleavage buffer and 3 nM DNA plasmid was added. After 10 min, the reaction was stopped by adding 6X loading dye (NEB) with EDTA and run 2% agarose gel. Similarly, gels were dyed with SYBR gold (ThermoFisher Scientific) and imaged with ChemiDoc XRS + System (Bio-Rad). Cleavage was quantified by ImageJ.

### *In vitro* cleavage assay for ssDNA and ssRNA

*In vitro* cleavage assay was performed with purified FCA anCas, BCA anCas and SpCas9 endonucleases. In all assays, 30 nM enzyme was incubated for 15 min with 30 nM sgRNA (Spy-sgRNA 20 nt) at 1:1 ratio in the cleavage buffer (100 mM NaCl, 50 mM Tris-HCl, 10 mM MgCl_2_, 100 µg/BSA, pH 7.9) at 37 °C. Then, 3 nM target (ssDNA or ssRNA) was added and incubated for different time intervals (0, 5, 10, 30 and 60 min). For ssDNA target, reaction was stopped by adding 6X loading dye (NEB) with urea. Samples were boiled for 10 min at 80 °C and were resolved by 2.5% denaturing urea agarose gel. For ssRNA target, reaction was stopped by adding 2X RNA gel-loading buffer (NEB) with urea. Samples were boiled for 10 min at 95 °C and were resolved by 15% denaturing urea polyacrylamide gel electrophoresis. In all cases, gels were dyed with SYBR gold (ThermoFisher Scientific) and imaged with ChemiDoc XRS + System (Bio-Rad). Cleavage was quantified by ImageJ and fitted with single-exponential decay curve.

### ELISA test

Elisa test was performed by using a modified protocol described elsewhere^60^. Briefly, 1 µg/well of SpCas9, FCA anCas, BCA anCas and bovine serum albumin (BSA, Sigma Aldrich) were diluted in 1x bicarbonate buffer and coated onto 96-well plates (ThermoFisher Scientific) overnight at 4 °C. Plates were washed with 1X wash buffer (TBST, ThermoFisher Scientific) and blocking with 1% BSA blocking solution for 1 hour at room temperature. Anti-Cas9 rabbit antibody (Rockland, 600-401-GK0) was diluted 1:25000 in 1% BSA blocking solution and plates were incubated for 2 hours at room temperature. Then, plates were washed and HRP-conjugated goat anti-Rabbit IgG (H+L) (Invitrogen), diluted 1:2000 in 1% BSA blocking solution, was added, and incubated for 1 hour at room temperature. Finally, 3,3′,5,5′-Tetramethylbenzidine ELISA substrate solution (ThermoFisher Scientific) was added and incubated for 10 min at room temperature. The reaction was stopped with 1 N sulfuric acid. The absorbance was measured at 450 nm by using a VICTOR X5 microplate reader (PerkinElmer).

### Human HEK293T cells *in vivo* cleavage

Functional validation of ancestral Cas nucleases was carried out in human HEK293T cells, as described elsewhere^61^. Cells were grown in DMEM medium (Dulbecco’ s Modified Eagle Medium, Gibco), supplemented with sterile-filtered 10% fetal bovine serum (FBS), 10 mM HEPES pH 7.4, 2 mM L-glutamine and penicillin (100 IU/ml)–streptomycin (100□μg/ml) and handled under aseptic conditions using a sterile hood. HEK293T cells were cultured in incubators at 37°C, 95% humidity and 5% CO_2_. Humanized anCas were cloned into pcDNA3.1 plasmid expression vector (ThermoFisher). The sgRNA target sequences were designed with Breaking-Cas web tool^62^ and cloned into MLM3636 plasmid vector (Addgene #43860) through Golden Gate cloning method. SpCas9 from hCas9 plasmid (Addgene #41815) was used as a positive control. For the *in vivo* genome-editing tests, cells were plated in 24-well plates at a density of 4 × 10^5^ cells/ml in a 0.5 ml volume of DMEM without antibiotics. To these cells, 1 μg of hCas/hanCas plasmid and 0.5 μg of the corresponding sgRNA plasmid were transfected with 2 μl of Lipofectamine 2000 (Life Technologies) diluted in 100 μl of Opti-MEM (Gibco) per well. 72 hours post-transfection genomic DNA was isolated with High Pure Template Preparation Kit (Roche). INDEL occurrence was assessed by T7 Endonuclease I assay on PCR-amplified DNA fragments surrounding the target DSB.

### Characterization of gene-edited allelic variants on TYR and OCA2 genes

DNA fragment covering the target region for the designed gRNAs at exon 1 of *TYR* (NM_000372.5; Gene ID: 7299) and exon 14 of *OCA2* (NM_000275.3; Gene ID: 4948), in HEK293T human cells was PCR-amplified using customized primers that included the adapters for NGS sequencing followed by the gene-specific sequence (Supplementary Information). Library preparation for all the samples in duplicate generated after cleavage and repair with humanized versions of SpCas9, PDCA, PCA, SCA, BCA and FCA anCas, was carried out as previously described^63^. Briefly, PCR products were indexed using the Nextera XT DNA library preparation kit (Illumina), purified, pooled, and sequenced in a MiSeq platform (Illumina, San Diego, CA, USA) using a 2×250 paired-end setting. The average depth of coverage was roughly 20.000x for each sample. Percentage of the different alleles after Cas9-mediated editing was determined using Mosaic Finder (MF)^42^. MF’s pipeline integrates NGS read mapping, normalization of read counts, mutation frequency calculation and genome-editing efficiency statistics at each position of the target region. MF takes, as input files, the fastq generated by pair-end sequencing and generates consensus sequences by joining the corresponding read pairs (forward and reverse). The repertoire of consensus sequences (allelic clusters) represents the allelic diversity generated by the Cas9-mediated edition. These clusters are then aligned against the sequence used as reference and are classified in allelic classes based on the different type of the identified mutation (mismatch, insertion/deletion). The frequency each cluster are then calculated and plotted. For multiple sequence alignment visualization of the different alleles classified by MF we have used the Jalview program (https://www.jalview.org/). A similar analysis was carried out using other available software such as CrispRVariants R-software package^43^ (data available upon request).

### Traffic Light reporter (TLR) cell generation

HEK293T cells were infected with TLR lentivirus at a 0.2 multiplicity of infection (MOI) to generate the TLR cell line as described before^64^. After infection, cells were selected with puromycin following standard protocols for 2 weeks before being used for subsequent assays. Cell transfection experiments were performed with Lipofectamine 3000 (Thermo Fisher Scientific) with 0.5 ug of nuclease and 0.5 ug of gRNA expression DNA vectors. Cells were analysed by flow cytometry using BD LSR Fortessa (BD Bioscience. Blue 488 nm laser with 530/30 filter and Yellow Green 561nm laser with 610/20 filter) 3-4 days after transfection. The relative NHEJ frequency was estimated by the number of RFP-positive cells. SpCas9 gRNA expression cassettes were cloned on TOPO vectors using manufacturing recommendations (Zero Blunt® TOPO®, Thermo Fisher Scientific).

## Data availability

We have made available sequencing data. Other data supporting the findings of this study are available from the corresponding authors upon reasonable request.

## Acknowledgements

This work has been supported by grants PID2019-109087RB-I00 to R.P.-J and grant RTI2018-101223-B-I00 to L.M. from Spanish Ministry of Science and Innovation. This project has received funding from the European Union’s Horizon 2020 research and innovation programme under grant agreement No 964764 to R.P.-J. The content presented in this document represents the views of the authors, and the European Commission has no liability in respect to the content. We acknowledge financial support from Spanish Foundation for the promotion of research of Amyotrophic Lateral Sclerosis (FUNDELA). A.F acknowledges CIBERER intramural funds (ER19P5AC756/2021). F.J.M.M acknowledges research support by Conselleria d’Educació, Investigació, Cultura i Esport from Generalitat Valenciana, research projects PROMETEO/2017/129 and PROMETEO/2021/057. M.M acknowledges funding from the Spanish Center for Biomedical Network Research on Rare Diseases (CIBERER) grant ER19P5AC728/2021. The work has received funding from the Regional Government of Madrid (CAM) grant B2017/ BMD3721 to M.A.M-P and from Instituto de Salud Carlos III cofounded with the European Regional Development Fund (ERDF), “A way to make Europe”) within the National Plans for Scientific and Technical Research and Innovation 2017–2020 and 2021–2024 (PI17/1659; PI20/0429 and IMP/00009) to M.A.M-P. B.P.K. was supported by an MGH ECOR Howard M. Goodman Award.

## Author contributions

R. P-J. conceived the project. R. P-J., B. A-L., B. P. K., A. S-M., M. G., F. J. M. M., M.A. M-P., L. M., designed research and planned experiments. R. P-J. performed the phylogenetic analysis and ancestral sequence reconstruction. B. A.-L., Y. J, S.S., cloned and expressed proteins and performed *in vitro* experiments. B. A.-L., Y. J, S.S., M.M, A. F., Y. B., S.F-P, L.S., A. S-M., M.G., M.A. M-P., and L. M, performed functional validation of anCas in mammalian cells, sequencing experiments and bioinformatic analysis. A-L, Y. J, R. P-J., J.A. G., A. R., A. D., and W. V., designed analysed and represented structural data. All authors participated in discussions and provided ideas for the work. R.P-J., F. J. M. M., M.A. M-P., L. M wrote the original paper and all authors revised and edited the manuscript.

## Additional information

Supplementary information: The online version contains supplementary material available at www.biorxiv.org.

Permissions information is available at www.biorxiv.org.

