## Supplementary material for "Evolution of CRISPR-associated Endonucleases as Inferred from Resurrected Proteins": Evolution of CRISPR_SI

^1^CIC nanoGUNE BRTA, San Sebastian, Spain.

^2^Servicio de Genética, Hospital Universitario Ramón y Cajal, IRYCIS and Centro de Investigaciones Biomédicas en Red de Enfermedades Raras (CIBERER), Madrid, Spain

^3^Department of Molecular and Cellular Biology, National Centre for Biotechnology (CNB-CSIC) and Centre for Biomedical Network Research on Rare Diseases (CIBERER-ISCIII), Madrid, Spain.

^4^INGEMM, Hospital Universitario La Paz, CIBERER-ISCIII, Madrid, Spain.

^5^Laboratorio de Estudios Cristalográficos, IACT (CSIC-UGR), Armilla, Granada, Spain

^6^Interuniversity Institute of Bioinformatics in Brussels, ULB-VUB, Brussels 1050, Belgium

^7^Structural Biology Brussels, Vrije Universiteit Brussel, Brussels 1050, Belgium

^8^Structural Biology Research Centre, VIB, Brussels 1050, Belgium.

^9^Center for Genomic Medicine and Department of Pathology, Massachusetts General Hospital, Boston, MA, 02114, USA

^10^Department of Pathology, Harvard Medical School, Boston, MA, 02115, USA

^11^Integra Therapeutics S.L., Barcelona, Spain.

^12^Department of Medicine and Life Sciences, Universitat Pompeu Fabra, Barcelona, Spain.

^13^Dpto. Fisiología, Genética y Microbiología and Instituto Multidisciplinar para el Estudio del Medio "Ramón Margalef”, Universidad de Alicante, Alicante, Spain.

^14^Ikerbasque Foundation for Science, Bilbao, Spain.

**Supplementary Tables**

| **Supplementary Table 1. Statistics analysis of pLDDT values obtained from AlphaFold2 structure prediction of anCas and SpCas9.** | | | | | |
| --- | --- | --- | --- | --- | --- |
|  | **Mean** | **SD** | **Min** | **Max** | **Num Values** |
| **FCA anCas** | 82.24 | 12.60 | 35.81 | 98.33 | 1340 |
| **BCA anCas** | 85.87 | 11.28 | 33.75 | 98.32 | 1340 |
| **SCA anCas** | 85.97 | 11.72 | 33.15 | 98.31 | 1368 |
| **PCA anCas** | 86.88 | 10.75 | 31.90 | 98.42 | 1368 |
| **PDCA anCas** | 87.87 | 10.27 | 36.52 | 98.58 | 1368 |
| **SpCas9** | 88.33 | 9.92 | 38.10 | 98.61 | 1368 |

| **Supplementary Table 2. RMSD values of different SpCas9 protein domains obtained from anCas.** | | | | | |
| --- | --- | --- | --- | --- | --- |
| **Domains** | **FCA anCas** | **BCA anCas** | **SCA anCas** | **PCA anCas** | **PDCA anCas** |
| **RuvCI** | 0.5820 | 0.3722 | 0.3414 | 0.3415 | 0.3617 |
| **BH** | 0.5877 | 0.3153 | 0.3453 | 0.3240 | 0.3272 |
| **REC1** | 0.8105 | 0.5504 | 0.6405 | 0.5821 | 0.4768 |
| **REC2** | 1.6140 | 0.8739 | 0.3722 | 0.5694 | 0.5194 |
| **REC** | 2.9309 | 1.1052 | 1.0937 | 0.9087 | 1.1426 |
| **RUVC II** | 0.8643 | 0.7294 | 0.6052 | 0.7951 | 0.7577 |
| **HNH** | 1.1783 | 0.8971 | 0.8710 | 0.9024 | 0.7063 |
| **RUVC III** | 2.1138 | 1.0338 | 1.0573 | 1.1097 | 0.9155 |
| **PI** | 1.9637 | 1.1284 | 1.0394 | 0.9443 | 0.7475 |

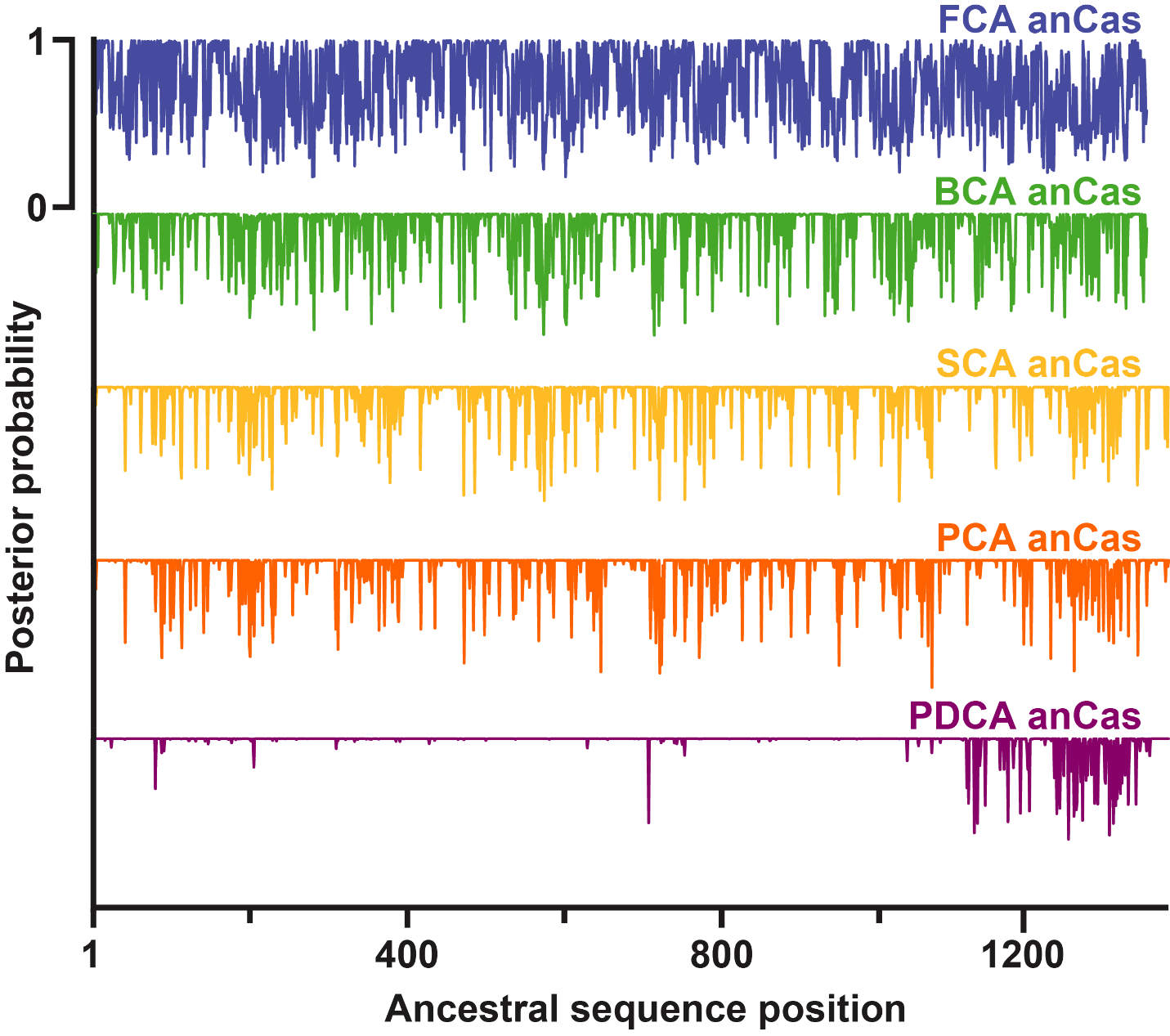

**Supplementary Figure 1. Posterior probability distribution for each inferred residue of all ancestral anCas endonucleases.** The residue with the highest posterior probability is assigned at each position. In all cases, posterior probability average is close to 1 except for FCA anCas which shows an average value of 0.74.

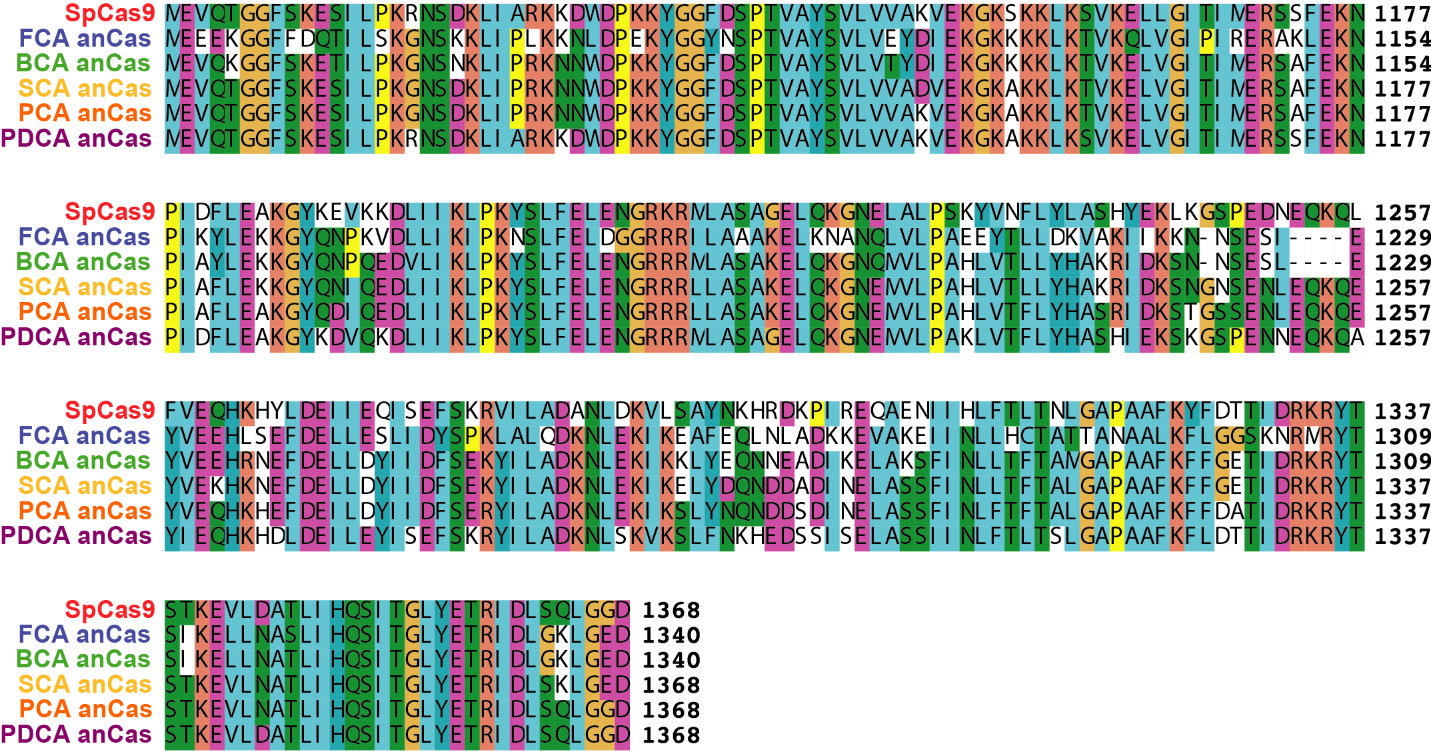

**Supplementary Figure 2. Alignment of the amino acid sequences from PI domain of anCas and SpCas9.**

| **Main residues involved in SpCas9 activity and their correspondence with anCas sequences. Mutations are highlighted in blue.** | | | | | |
| --- | --- | --- | --- | --- | --- |
| **SpCas9** | **FCA anCas** | **BCA anCas** | **SCA anCas** | **PCA anCas** | **PDCA anCas** |
| D10 | D10 | D10 | D10 | D10 | D10 |
| S15 | S15 | S15 | S15 | S15 | S15 |
| R66 | R66 | R66 | R66 | R66 | R66 |
| R70 | R70 | R70 | R70 | R70 | R70 |
| R74 | R74 | R74 | R74 | R74 | R74 |
| R78 | R78 | R78 | R78 | R78 | R78 |
| P475 | P475 | P475 | P475 | P475 | P475 |
| W476 | W476 | W476 | W476 | W476 | W476 |
| N477 | N477 | N477 | N477 | N477 | N477 |
| H840 | H838 | H838 | H840 | H840 | H840 |
| K1107 | D1084 | K1084 | K1107 | K1107 | K1107 |
| S1109 | T1086 | T1086 | S1109 | S1109 | S1109 |
| W1126 | L1103 | W1103 | W1126 | W1126 | W1126 |
| R1333 | R1305 | R1305 | R1333 | R1333 | R1333 |
| K1334 | M1306 | K1306 | K1334 | K1334 | K1334 |
| R1335 | R1307 | R1307 | R1335 | R1335 | R1335 |

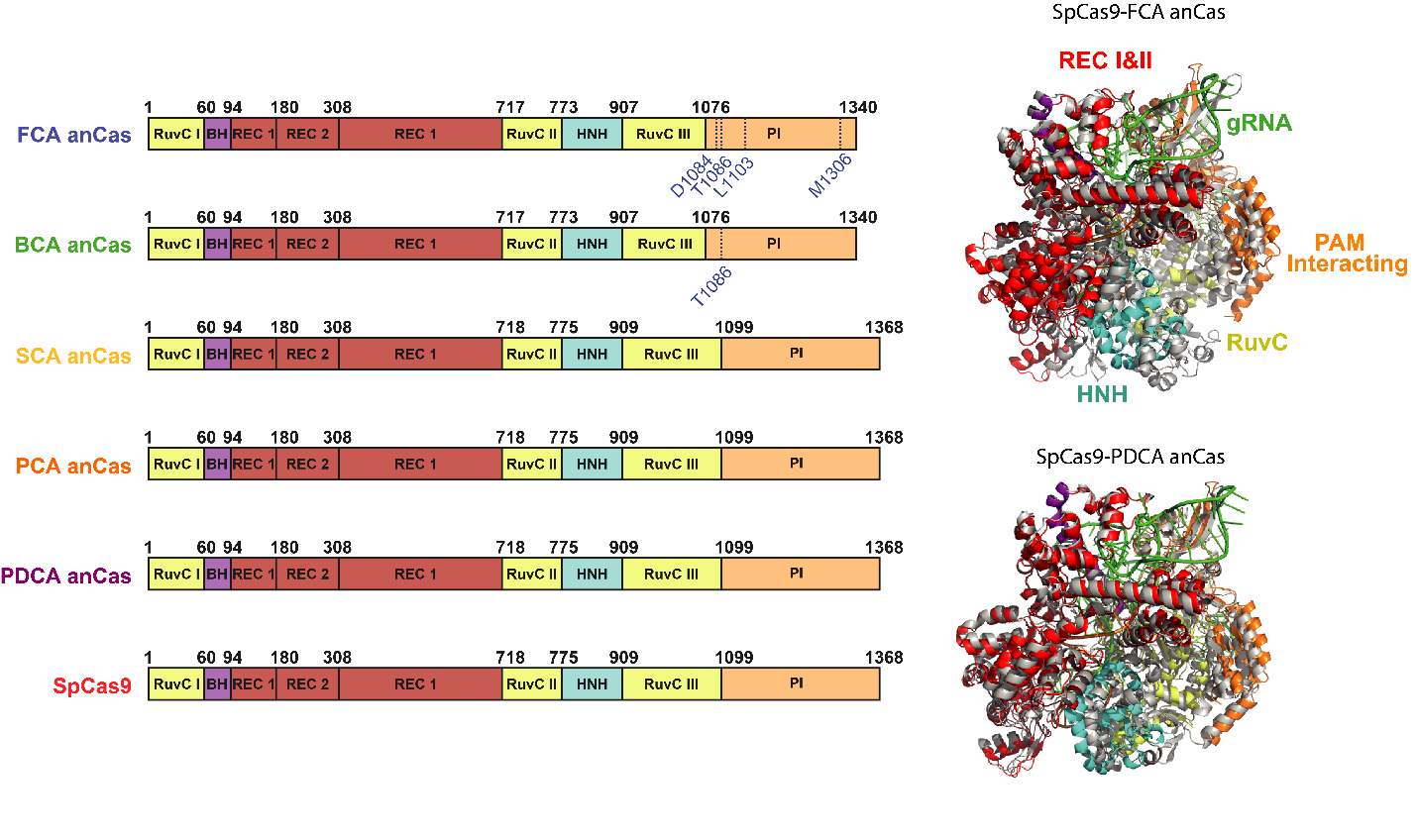

**Supplementary Figure 3. List of important mutations and domain organization of anCas compared to SpCas9.** Mutations of the main residues involved in PAM recognition are marked in blue. Bottom figure depicts domain organization and structural alignment of SpCas9-FCA and SpCas9 PDCA anCas. Ancestral anCas are grey colored and SpCas9 colored by domains.

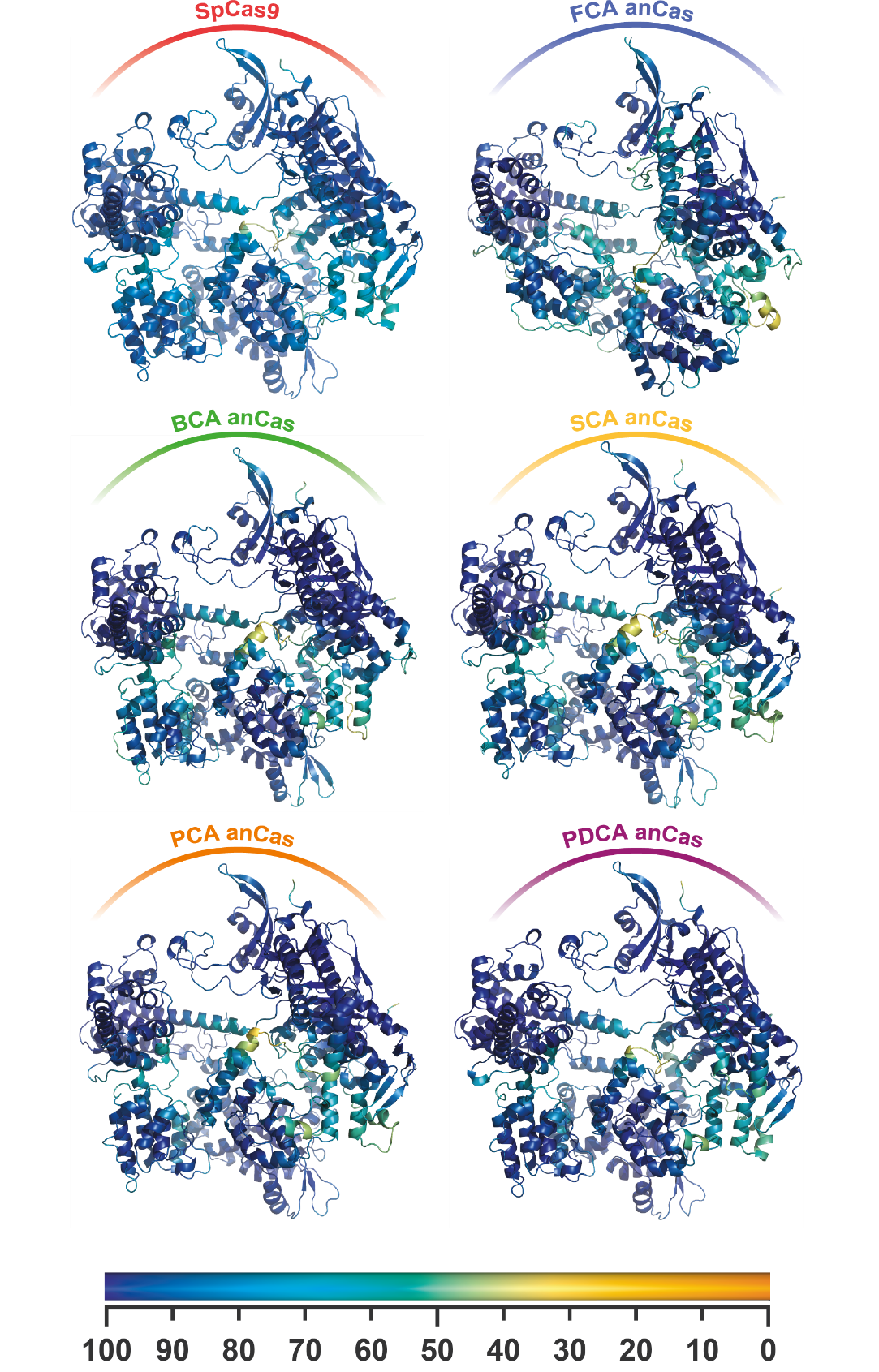

**Supplementary Figure 4. Structural predictions of anCas and SpCas9 by AlphaFold2.** Structures are colored by pLDDT score according to the color bar.

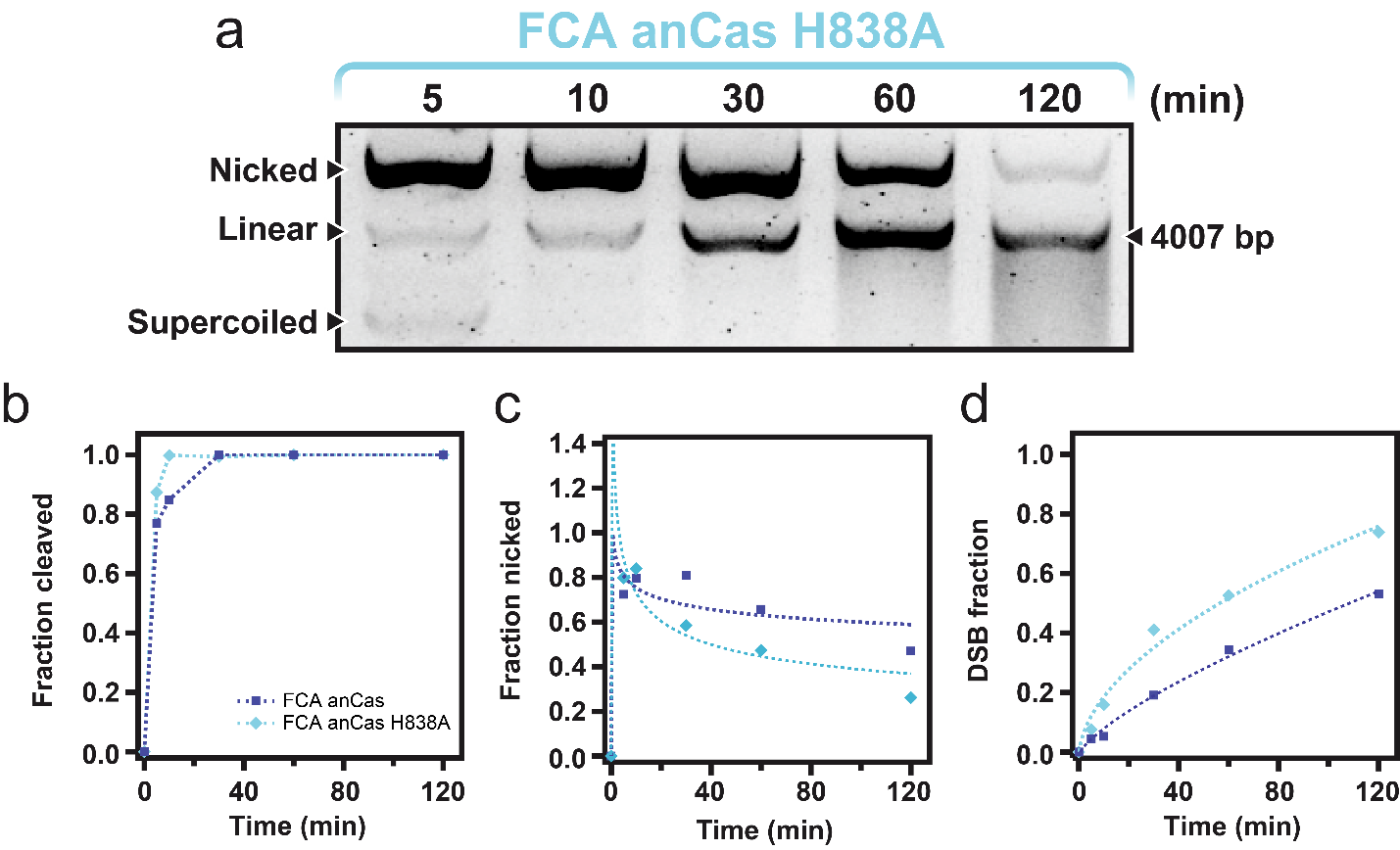

| **Kinetics parameters of the FCA anCas and FCA anCas H838A determined from the single exponential decay curves.** | | | | |
| --- | --- | --- | --- | --- |
|  | **FCA anCas** | | **FCA anCas H838A** | |
| **Kinetics parameters** | Total | DSB | Total | DSB |
| *K_cleave_* (min^-1^) | 0.27 ± 0.04 | 0.009 ± 0.002 | 0.33 ± 0.03 | 0.021 ± 0.004 |
| Max fraction | 1 | 0.530 | 1 | 0.73 |
| R^2^ | 0.9906 | 0.9982 | 0.9986 | 0.9917 |

**Supplementary Figure 5. Activity of FCA anCas H838A endonuclease on a supercoiled DNA substrate (a)** *In vitro* cleavage assay for anCas FCA H838A on a 4007 bp substrate at different reaction times showing nicked and linear fractions. **(b)** Quantification of total cleavage fraction at different reaction times and exponential fits (lines). **(c)** Quantification of fraction nicked at different times. **(d)** Quantification of DSB cleavage. Single-exponential fits were used to obtain *k*_cleave_ and maximum fraction cleaved (amplitude).

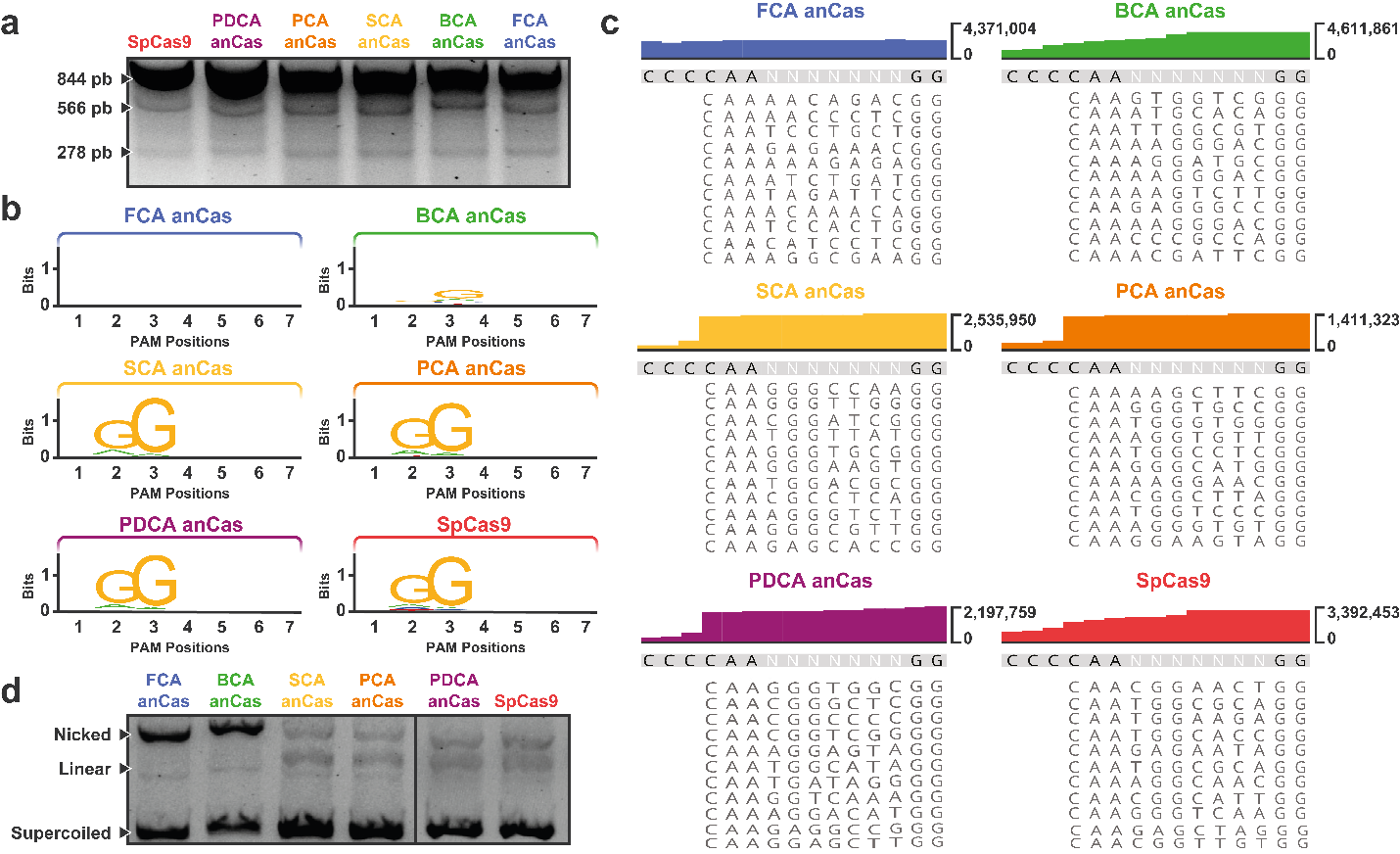

**Supplementary Figure 6. PAM determination of anCas. (a)** Example of *in vitro* cleavage assay to obtain 278 bp fragment for NGS analysis. **(b)** Weblogo of the different PAM recognized by anCas and SpCas9. **(c)** Reads of 278 bp fragments analyzed by NGS for PAM determination. Y-axis corresponds to the total mapped reads and X-axis corresponds to bp analyzed (examples of different sequences obtained are shown). **(d)** *In vitro* cleavage assay using the PAM sequence TCC.

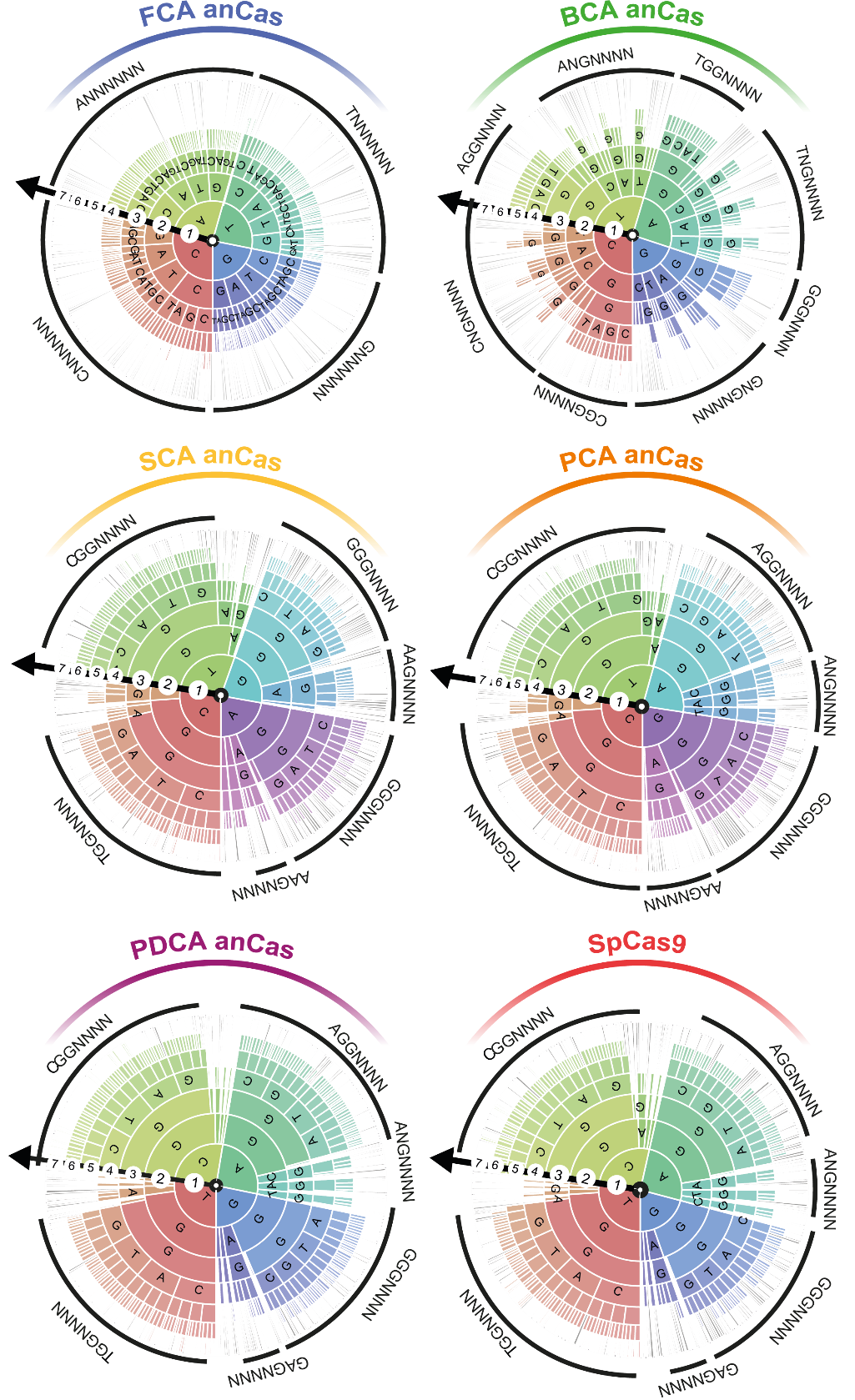

**Supplementary Figure 7.** **PAM wheels (Krona plots) for all five anCas and SpCas9 including 7-nucleotides PAM analysis.** A preference for NGG PAM is observed except for FCA anCas.

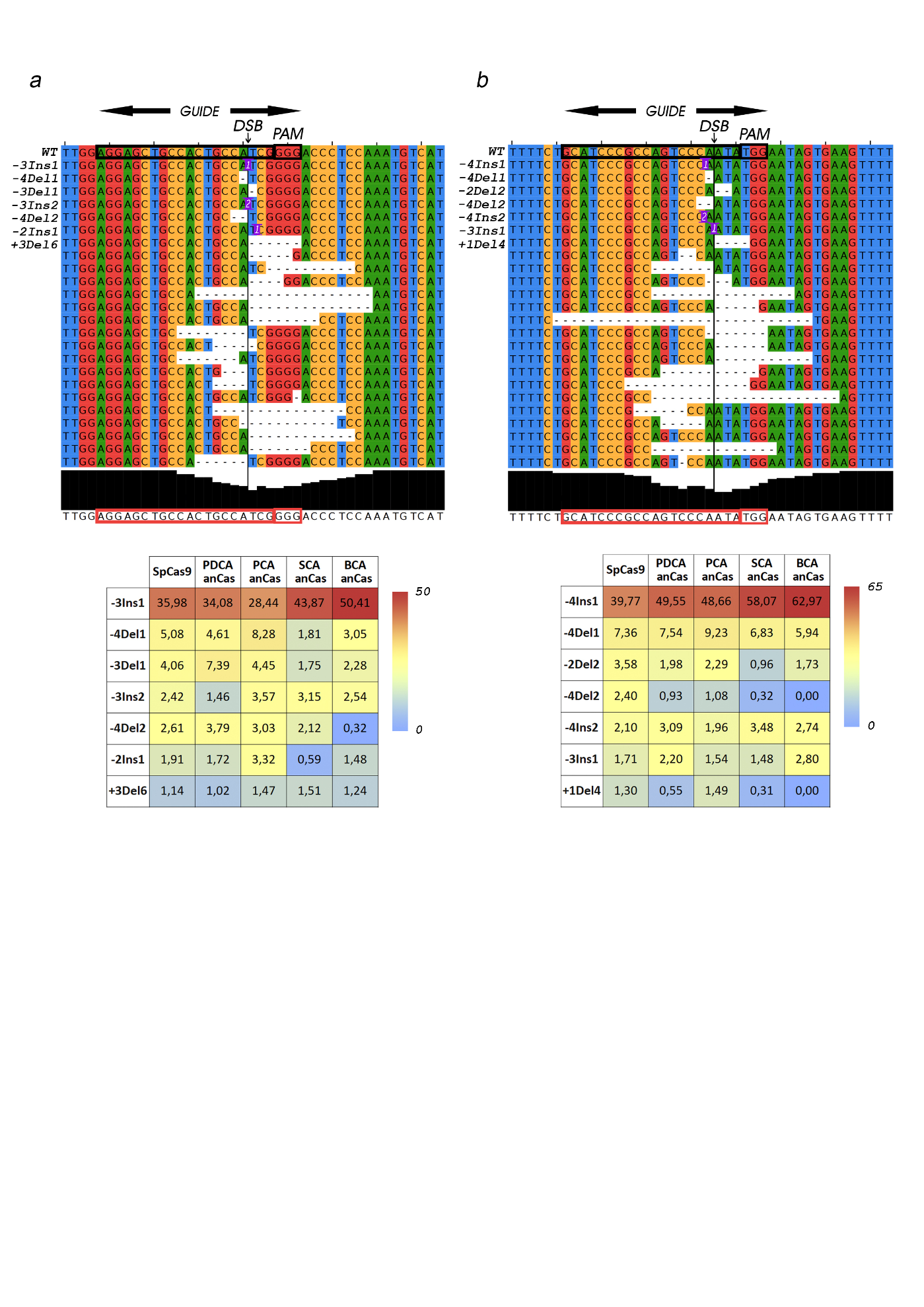

**Supplementary Figure 8. Analysis of the *in vivo* activity of anCas variants*.***Alignments generated by Jalview program of the wild-type and the most frequent edited alleles (indels) detected by Mosaic Finder in **(a)** *OCA2* and **(b)** *TYR* genes after NHEJ cell repair in HEK293T cells. Heatmaps are shown underneath the alignments highlighting the frequencies of the top-7 most frequent alleles generated after cleavage and repair with SpCas9, PDCA, PCA, SCA and BCA anCas, once normalized with respect to the total number of indels for each Cas. The guide, the PAM and the DSB theoretical site are marked in the figure. For the mutation nomenclature of each allele we consider the first nucleotide of the PAM as +1. Numbers within the allele sequences represent the length of insertions or deletion in the exact location indicated by the first figure. Example: *-4Ins1*, insertion of 1 nucleotide four bases upstream the PAM.

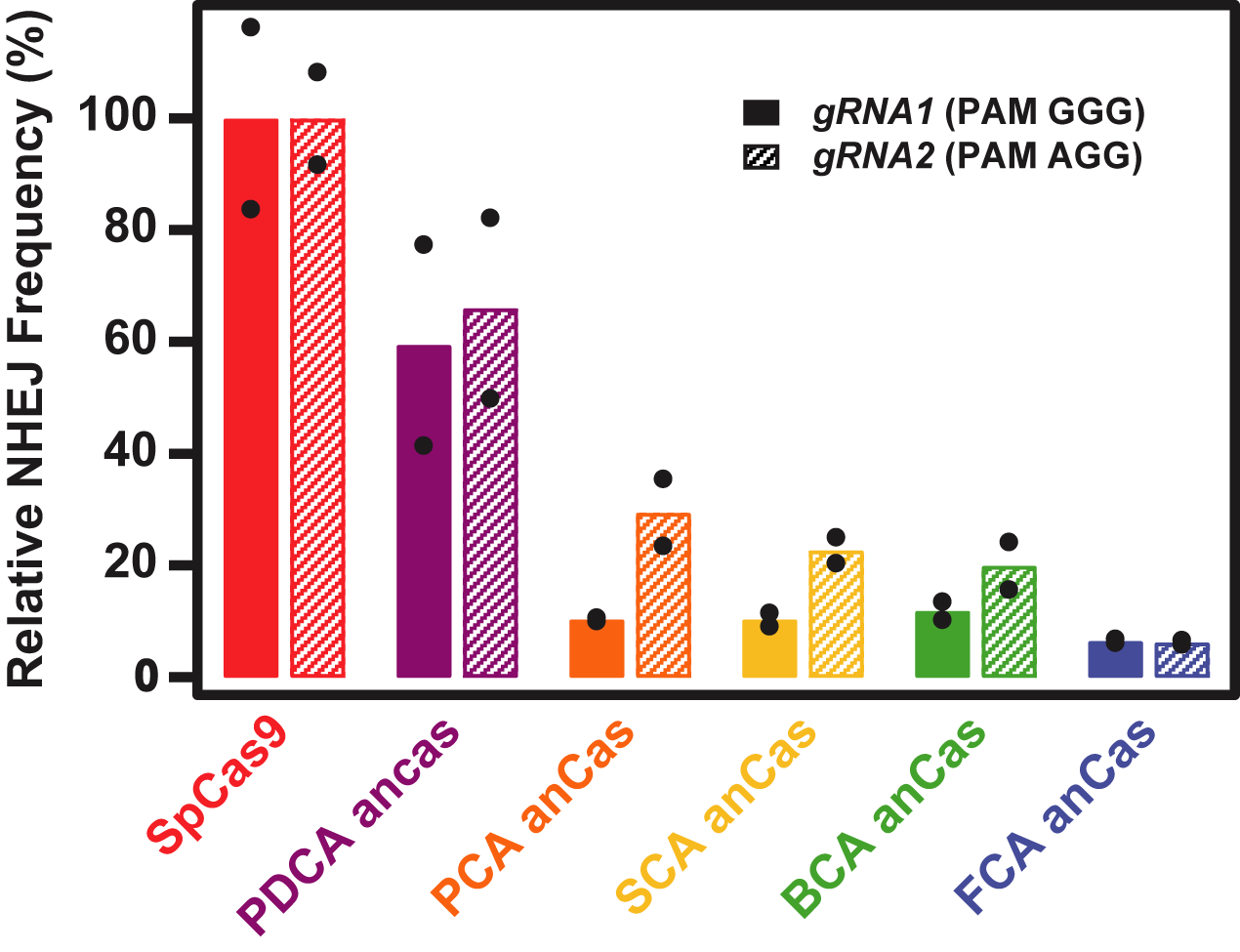

**Supplementary Figure 9.** **Traffic Light Reporter cleavage assay targeting gene *TLR***. The relative NHEJ frequency is estimated by the number of RFP-positive cells and is normalized to SpCas9.

**Sequences Information**

| **List of Cas9 Sequences utilized for Ancestral Reconstruction of ancient endonucleases.** | | | | |
| --- | --- | --- | --- | --- |
| **Phylum** | **Class** | **Genus** | **Specie** | **Sequence** |
| Firmicutes | Bacilli | Streptoctoccus | *Streptococcus dysgalactiae* | [WP_084916602.1](https://www.ncbi.nlm.nih.gov/protein/WP_084916602.1) |
| Firmicutes | Bacilli | Streptoctoccus | *Streptococcus canis* | [WP_003043819.1](https://www.ncbi.nlm.nih.gov/protein/WP_003043819.1) |
| Firmicutes | Bacilli | Streptoctoccus | *Streptococcus gallolyticus* | [WP_012962174.1](https://www.ncbi.nlm.nih.gov/protein/WP_012962174.1) |
| Firmicutes | Bacilli | Streptoctoccus | *Streptococcus infantarius* | [WP_014334983.1](https://www.ncbi.nlm.nih.gov/protein/WP_014334983.1) |
| Firmicutes | Bacilli | Streptoctoccus | *Streptococcus phocae* | [WP_054279288.1](https://www.ncbi.nlm.nih.gov/protein/WP_054279288.1) |
| Firmicutes | Bacilli | Streptoctoccus | *Streptococcus equinus* | [WP_157339387.1](https://www.ncbi.nlm.nih.gov/protein/WP_157339387.1) |
| Firmicutes | Bacilli | Streptoctoccus | *Streptococcus pasteurianus* | [WP_061100419](https://www.ncbi.nlm.nih.gov/protein/WP_061100419) |
| Firmicutes | Bacilli | Streptoctoccus | *Streptococcus equi* | [WP_037581760.1](https://www.ncbi.nlm.nih.gov/protein/WP_037581760.1?report=genpept) |
| Firmicutes | Bacilli | Streptoctoccus | *Streptococcus mutans* | [WP_002279859.1](https://www.ncbi.nlm.nih.gov/protein/WP_002279859.1) |
| Firmicutes | Bacilli | Streptoctoccus | *Streptococcus iniae* | [WP_003099269.1](https://www.ncbi.nlm.nih.gov/protein/WP_003099269.1) |
| Firmicutes | Bacilli | Streptoctoccus | *Streptococcus agalactiae* | [AFV72233.1](https://www.ncbi.nlm.nih.gov/protein/AFV72233.1) |
| Firmicutes | Bacilli | Streptoctoccus | *Streptococcus caballi* | [WP_018363470.1](https://www.ncbi.nlm.nih.gov/protein/WP_018363470.1) |
| Firmicutes | Bacilli | Streptoctoccus | *Streptococcus anginosus* | [WP_003041502](https://www.ncbi.nlm.nih.gov/protein/WP_003041502) |
| Firmicutes | Bacilli | Streptoctoccus | *Streptococcus varani* | [WP_093650272](https://www.ncbi.nlm.nih.gov/protein/WP_093650272) |
| Firmicutes | Bacilli | Streptoctoccus | *Streptococcus intermedius* | [WP_082312238.1](https://www.ncbi.nlm.nih.gov/protein/WP_082312238.1) |
| Firmicutes | Bacilli | Streptoctoccus | *Streptococcus ratti* | [WP_003088697. 1](https://www.ncbi.nlm.nih.gov/protein/489179199) |
| Firmicutes | Bacilli | Streptoctoccus | *Streptococcus sanguinis* | [WP_002906454.1](https://www.ncbi.nlm.nih.gov/protein/WP_002906454.1) |
| Firmicutes | Bacilli | Streptoctoccus | *Streptococcus mitis* | [WP_084927115.1](https://www.ncbi.nlm.nih.gov/protein/WP_084927115.1) |
| Firmicutes | Bacilli | Streptoctoccus | *Streptococcus gordonii* | [WP_045635197.1](https://www.ncbi.nlm.nih.gov/protein/WP_045635197.1) |
| Firmicutes | Bacilli | Streptoctoccus | *Streptococcus suis* | [WP_044681799.1](https://www.ncbi.nlm.nih.gov/protein/WP_044681799.1) |
| Firmicutes | Bacilli | Streptoctoccus | *Streptococcus salivarius* | [WP_002891502.1](https://www.ncbi.nlm.nih.gov/protein/WP_002891502.1) |
| Firmicutes | Bacilli | Streptoctoccus | *Streptococcus pseudoporcinus* | [WP_007896501.1](https://www.ncbi.nlm.nih.gov/protein/WP_007896501.1) |
| Firmicutes | Bacilli | Streptoctoccus | *Streptococcus henryi* | [WP_074484960.1](https://www.ncbi.nlm.nih.gov/protein/WP_074484960.1) |
| Firmicutes | Bacilli | Streptoctoccus | *Streptococcus thermophilus* | [WP_065972475.1](https://www.ncbi.nlm.nih.gov/protein/WP_065972475.1) |
| Firmicutes | Bacilli | Streptoctoccus | *Streptococcus parauberis* | [WP_076751909.1](https://www.ncbi.nlm.nih.gov/protein/WP_076751909.1) |
| Firmicutes | Bacilli | Streptoctoccus | *Streptococcus constellatus* | [WP_006269658.1](https://www.ncbi.nlm.nih.gov/protein/WP_006269658.1) |
| Firmicutes | Bacilli | Streptoctoccus | *Streptococcus pyogenes* | [WP_032464890.1](https://www.ncbi.nlm.nih.gov/protein/WP_032464890.1) |
| Firmicutes | Bacilli | Streptoctoccus | *Streptococcus cuniculi* | [WP_075103982.1](https://www.ncbi.nlm.nih.gov/protein/WP_075103982.1) |
| Firmicutes | Bacilli | Enterococcus | *Enterococcus italicus* | [WP_007209003.1](https://www.ncbi.nlm.nih.gov/protein/WP_007209003.1) |
| Firmicutes | Bacilli | Enterococcus | *Enterococcus massiliensis* | [WP_048604708.1](https://www.ncbi.nlm.nih.gov/protein/WP_048604708.1) |
| Firmicutes | Bacilli | Enterococcus | *Enterococcus phoeniculicola* | [OJG69383.1](https://www.ncbi.nlm.nih.gov/protein/OJG69383.1) |
| Firmicutes | Bacilli | Enterococcus | *Enterococcus faecalis* | [QKR88578.1](https://www.ncbi.nlm.nih.gov/protein/QKR88578.1) |
| Firmicutes | Bacilli | Enterococcus | *Enterococcus devriesei* | [WP_071862060.1](https://www.ncbi.nlm.nih.gov/protein/WP_071862060.1) |
| Firmicutes | Bacilli | Enterococcus | *Enterococcus hirae* | [WP_096708680.1](https://www.ncbi.nlm.nih.gov/protein/1248210339) |
| Firmicutes | Bacilli | Enterococcus | *Enterococcus faecium* | [WP_087063969.1](https://www.ncbi.nlm.nih.gov/protein/WP_087063969.1) |
| Firmicutes | Bacilli | Enterococcus | *Enterococcus pseudoavium* | [WP_067627992.1](https://www.ncbi.nlm.nih.gov/protein/WP_067627992.1) |
| Firmicutes | Bacilli | Enterococcus | *Enterococcus durans* | [WP_081134165.1](https://www.ncbi.nlm.nih.gov/protein/WP_081134165.1) |
| Firmicutes | Bacilli | Enterococcus | *Enterococcus mundtii* | [WP 023519017.1](https://www.ncbi.nlm.nih.gov/protein/WP_023519017.1) |
| Firmicutes | Bacilli | Vagococcus | *Vagococcus fluvialis* | [WP_207053108.1](https://www.ncbi.nlm.nih.gov/protein/WP_207053108.1) |
| Firmicutes | Bacilli | Vagococcus | *Vagococcus teuberi* | [WP 071456514.1](https://www.ncbi.nlm.nih.gov/protein/WP_071456514.1) |
| Firmicutes | Bacilli | Listeria | *Listeria monocytogenes* | [WP_003739838.1](https://www.ncbi.nlm.nih.gov/protein/WP_003739838.1) |
| Firmicutes | Bacilli | Listeria | *Listeria ivanovii* | [WP_038409211.1](https://www.ncbi.nlm.nih.gov/protein/WP_038409211.1) |
| Firmicutes | Bacilli | Listeria | *Listeria innocua* | [WP_010991369.1](https://www.ncbi.nlm.nih.gov/protein/WP_010991369.1) |
| Firmicutes | Bacilli | Listeria | *Listeria seeligeri* | [WP_075702521.1](https://www.ncbi.nlm.nih.gov/protein/WP_075702521.1) |
| Firmicutes | Bacilli | Halolactibacillus | *Halolactibacillus alkaliphilus* | [WP_089800158.1](https://www.ncbi.nlm.nih.gov/protein/WP_089800158.1) |
| Firmicutes | Bacilli | Pelagirhabdus | *Pelagirhabdus alkalitolerans* | [WP_090793453.1](https://www.ncbi.nlm.nih.gov/protein/WP_090793453.1) |
| Firmicutes | Bacilli | Dolosigranulum | *Dolosigranulum pigrum* | [WP_004636532.1](https://www.ncbi.nlm.nih.gov/protein/WP_004636532.1) |
| Firmicutes | Bacilli | Anaerostipes | *Anaerostipes hadrus* | [WP_173774801.1](https://www.ncbi.nlm.nih.gov/protein/WP_173774801.1) |
| Firmicutes | Bacilli | Floricoccus | *Floricoccus tropicus* | [WP_070791099.1](https://www.ncbi.nlm.nih.gov/protein/WP_070791099.1) |
| Firmicutes | Bacilli | Urinacoccus | *Urinacoccus massiliensis* | [WP_034440723](https://www.ncbi.nlm.nih.gov/protein/WP_034440723) |
| Firmicutes | Clostridia | Clostridium | *Clostridium sp CAG299* | [CDD37961.1](https://www.ncbi.nlm.nih.gov/protein/CDD37961.1) |
| Firmicutes | Clostridia | Clostridium | *Clostridium sp CAG964* | [CDC80610.1](https://www.ncbi.nlm.nih.gov/protein/CDC80610.1) |
| Firmicutes | Clostridia | Clostridium | *Clostridium_sp_CAG122* | [CCZ42109.1](https://www.ncbi.nlm.nih.gov/protein/CCZ42109.1) |
| Firmicutes | Clostridia | Lachnospira | *Lachnospira multipara* | [WP_027438114.1](https://www.ncbi.nlm.nih.gov/protein/WP_027438114.1) |
| Firmicutes | Clostridia | Ruminococcus | *Ruminococcus lactaris* | [WP_005609677.1](https://www.ncbi.nlm.nih.gov/protein/WP_005609677.1) |
| Firmicutes | Clostridia | Dorea | *Dorea longicatena* | [WP_055214841.1](https://www.ncbi.nlm.nih.gov/protein/WP_055214841.1) |
| Actinobacteria | Coriobacteriaceae | Olsonella | *Olsenella_DNF00959* | [WP_062531800.1](https://www.ncbi.nlm.nih.gov/protein/WP_062531800.1) |
| Actinobacteria | Coriobacteriaceae | Olsonella | *Olsenella profusa* | [WP_021725096.1](https://www.ncbi.nlm.nih.gov/protein/WP_021725096.1) |
| Actinobacteria | Bifidobacteriaceae | Bifidobacterium | *Bifidobacterium pseudocatenulatum* | [WP_065439263.1](https://www.ncbi.nlm.nih.gov/protein/WP_065439263.1) |

**PAM library cloned in PUC18 (Figure 3)**

Target sequence is indicated in blue and random PAM in red

tcgcgcgtttcggtgatgacggtgaaaacctctgacacatgcagctcccggagacggtcacagcttgtctgtaagcggatgccgggagcagacaagcccgtcagggcgcgtcagcgggtgttggcgggtgtcggggctggcttaactatgcggcatcagagcagattgtactgagagtgcaccatatgcggtgtgaaataccgcacagatgcgtaaggagaaaataccgcatcaggcgccattcgccattcaggctgcgcaactgttgggaagggcgatcggtgcgggcctcttcgctattacgccagctggcgaaagggggatgtgctgcaaggcgattaagttgggtaacgccagggttttcccagtcacgacgttgtaaaacgacggccagtgccaagcttgcatgcctgcaggtcgactctagagggatccagcaacaacggtcggccacaccttccattgtcgtggccacgctcggattacacggcagaggtgcttgtgttccgacaggctagcatattgtcctaaggcgttaccccaaNNNNNNNggtaccgagctcgaattcgtaatcatggtcatagctgtttcctgtgtgaaattgttatccgctcacaattccacacaacatacgagccggaagcataaagtgtaaagcctggggtgcctaatgagtgagctaactcacattaattgcgttgcgctcactgcccgctttccagtcgggaaacctgtcgtgccagctgcattaatgaatcggccaacgcgcggggagaggcggtttgcgtattgggcgctcttccgcttcctcgctcactgactcgctgcgctcggtcgttcggctgcggcgagcggtatcagctcactcaaaggcggtaatacggttatccacagaatcaggggataacgcaggaaagaacatgtgagcaaaaggccagcaaaaggccaggaaccgtaaaaaggccgcgttgctggcgtttttccataggctccgcccccctgacgagcatcacaaaaatcgacgctcaagtcagaggtggcgaaacccgacaggactataaagataccaggcgtttccccctggaagctccctcgtgcgctctcctgttccgaccctgccgcttaccggatacctgtccgcctttctcccttcgggaagcgtggcgctttctcaaagctcacgctgtaggtatctcagttcggtgtaggtcgttcgctccaagctgggctgtgtgcacgaaccccccgttcagcccgaccgctgcgccttatccggtaactatcgtcttgagtccaacccggtaagacacgacttatcgccactggcagcagccactggtaacaggattagcagagcgaggtatgtaggcggtgctacagagttcttgaagtggtggcctaactacggctacactagaagaacagtatttggtatctgcgctctgctgaagccagttaccttcggaaaaagagttggtagctcttgatccggcaaacaaaccaccgctggtagcggtggtttttttgtttgcaagcagcagattacgcgcagaaaaaaaggatctcaagaagatcctttgatcttttctacggggtctgacgctcagtggaacgaaaactcacgttaagggattttggtcatgagattatcaaaaaggatcttcacctagatccttttaaattaaaaatgaagttttaaatcaatctaaagtatatatgagtaaacttggtctgacagttaccaatgcttaatcagtgaggcacctatctcagcgatctgtctatttcgttcatccatagttgcctgactccccgtcgtgtagataactacgatacgggagggcttaccatctggccccagtgctgcaatgataccgcgagacccacgctcaccggctccagatttatcagcaataaaccagccagccggaagggccgagcgcagaagtggtcctgcaactttatccgcctccatccagtctattaattgttgccgggaagctagagtaagtagttcgccagttaatagtttgcgcaacgttgttgccattgctacaggcatcgtggtgtcacgctcgtcgtttggtatggcttcattcagctccggttcccaacgatcaaggcgagttacatgatcccccatgttgtgcaaaaaagcggttagctccttcggtcctccgatcgttgtcagaagtaagttggccgcagtgttatcactcatggttatggcagcactgcataattctcttactgtcatgccatccgtaagatgcttttctgtgactggtgagtactcaaccaagtcattctgagaatagtgtatgcggcgaccgagttgctcttgcccggcgtcaatacgggataataccgcgccacatagcagaactttaaaagtgctcatcattggaaaacgttcttcggggcgaaaactctcaaggatcttaccgctgttgagatccagttcgatgtaacccactcgtgcacccaactgatcttcagcatcttttactttcaccagcgtttctgggtgagcaaaaacaggaaggcaaaatgccgcaaaaaagggaataagggcgacacggaaatgttgaatactcatactcttcctttttcaatattattgaagcatttatcagggttattgtctcatgagcggatacatatttgaatgtatttagaaaaataaacaaataggggttccgcgcacatttccccgaaaagtgccacctgacgtctaagaaaccattattatcatgacattaacctataaaaataggcgtatcacgaggccctttcgtc

***In vitro* cleavage PAM sequences (Figure 3)**

Target sequence indicated in blue and PAM sequence in red

| **PAM** | **Sequence** |
| --- | --- |
| **TAC** | tgtgaaataccgcacagatgcgtaaggagaaaataccgcatcaggcgccattcgccattcaggctgcgcaactgttgggaagggcgatcggtgcgggcctcttcgctattacgccagctggcgaaagggggatgtgctgcaaggcgattaagttgggtaacgccagggttttcccagtcacgacgttgtaaaacgacggccagtgccaagcttgcatgcctgcaggtcgactctagagggatccagcaacaacggtcggccacaccttccattgtcgtggccacgctcggattacacggcagaggtgcttgtgttccgacaggctagcatattgtcctaaggcgttaccccaa**tac**gaggggtaccgagctcgaattcgtaatcatggtcatagctgtttcctgtgtgaaattgttatccgctcacaattccacacaacatacgagccggaagcataaagtgtaaagcctggggtgcctaatgagtga |
| **TGG** | tgtgaaataccgcacagatgcgtaaggagaaaataccgcatcaggcgccattcgccattcaggctgcgcaactgttgggaagggcgatcggtgcgggcctcttcgctattacgccagctggcgaaagggggatgtgctgcaaggcgattaagttgggtaacgccagggttttcccagtcacgacgttgtaaaacgacggccagtgccaagcttgcatgcctgcaggtcgactctagagggatccagcaacaacggtcggccacaccttccattgtcgtggccacgctcggattacacggcagaggtgcttgtgttccgacaggctagcatattgtcctaaggcgttaccccaa**tgg**gaggggtaccgagctcgaattcgtaatcatggtcatagctgtttcctgtgtgaaattgttatccgctcacaattccacacaacatacgagccggaagcataaagtgtaaagcctggggtgcctaatgagtga |
| **TAT** | tgtgaaataccgcacagatgcgtaaggagaaaataccgcatcaggcgccattcgccattcaggctgcgcaactgttgggaagggcgatcggtgcgggcctcttcgctattacgccagctggcgaaagggggatgtgctgcaaggcgattaagttgggtaacgccagggttttcccagtcacgacgttgtaaaacgacggccagtgccaagcttgcatgcctgcaggtcgactctagagggatccagcaacaacggtcggccacaccttccattgtcgtggccacgctcggattacacggcagaggtgcttgtgttccgacaggctagcatattgtcctaaggcgttaccccaa**tat**gaggggtaccgagctcgaattcgtaatcatggtcatagctgtttcctgtgtgaaattgttatccgctcacaattccacacaacatacgagccggaagcataaagtgtaaagcctggggtgcctaatgagtga |
| **TCC** | tgtgaaataccgcacagatgcgtaaggagaaaataccgcatcaggcgccattcgccattcaggctgcgcaactgttgggaagggcgatcggtgcgggcctcttcgctattacgccagctggcgaaagggggatgtgctgcaaggcgattaagttgggtaacgccagggttttcccagtcacgacgttgtaaaacgacggccagtgccaagcttgcatgcctgcaggtcgactctagagggatccagcaacaacggtcggccacaccttccattgtcgtggccacgctcggattacacggcagaggtgcttgtgttccgacaggctagcatattgtcctaaggcgttaccccaa**tcc**gaggggtaccgagctcgaattcgtaatcatggtcatagctgtttcctgtgtgaaattgttatccgctcacaattccacacaacatacgagccggaagcataaagtgtaaagcctggggtgcctaatgagtga |
| **CCC** | tgtgaaataccgcacagatgcgtaaggagaaaataccgcatcaggcgccattcgccattcaggctgcgcaactgttgggaagggcgatcggtgcgggcctcttcgctattacgccagctggcgaaagggggatgtgctgcaaggcgattaagttgggtaacgccagggttttcccagtcacgacgttgtaaaacgacggccagtgccaagcttgcatgcctgcaggtcgactctagagggatccagcaacaacggtcggccacaccttccattgtcgtggccacgctcggattacacggcagaggtgcttgtgttccgacaggctagcatattgtcctaaggcgttaccccaa**ccc**gaggggtaccgagctcgaattcgtaatcatggtcatagctgtttcctgtgtgaaattgttatccgctcacaattccacacaacatacgagccggaagcataaagtgtaaagcctggggtgcctaatgagtga |
| **TTT** | tgtgaaataccgcacagatgcgtaaggagaaaataccgcatcaggcgccattcgccattcaggctgcgcaactgttgggaagggcgatcggtgcgggcctcttcgctattacgccagctggcgaaagggggatgtgctgcaaggcgattaagttgggtaacgccagggttttcccagtcacgacgttgtaaaacgacggccagtgccaagcttgcatgcctgcaggtcgactctagagggatccagcaacaacggtcggccacaccttccattgtcgtggccacgctcggattacacggcagaggtgcttgtgttccgacaggctagcatattgtcctaaggcgttaccccaa**ttt**gaggggtaccgagctcgaattcgtaatcatggtcatagctgtttcctgtgtgaaattgttatccgctcacaattccacacaacatacgagccggaagcataaagtgtaaagcctggggtgcctaatgagtga |
| **TTC** | tgtgaaataccgcacagatgcgtaaggagaaaataccgcatcaggcgccattcgccattcaggctgcgcaactgttgggaagggcgatcggtgcgggcctcttcgctattacgccagctggcgaaagggggatgtgctgcaaggcgattaagttgggtaacgccagggttttcccagtcacgacgttgtaaaacgacggccagtgccaagcttgcatgcctgcaggtcgactctagagggatccagcaacaacggtcggccacaccttccattgtcgtggccacgctcggattacacggcagaggtgcttgtgttccgacaggctagcatattgtcctaaggcgttaccccaa**ttc**gaggggtaccgagctcgaattcgtaatcatggtcatagctgtttcctgtgtgaaattgttatccgctcacaattccacacaacatacgagccggaagcataaagtgtaaagcctggggtgcctaatgagtga |
| **TCA** | tgtgaaataccgcacagatgcgtaaggagaaaataccgcatcaggcgccattcgccattcaggctgcgcaactgttgggaagggcgatcggtgcgggcctcttcgctattacgccagctggcgaaagggggatgtgctgcaaggcgattaagttgggtaacgccagggttttcccagtcacgacgttgtaaaacgacggccagtgccaagcttgcatgcctgcaggtcgactctagagggatccagcaacaacggtcggccacaccttccattgtcgtggccacgctcggattacacggcagaggtgcttgtgttccgacaggctagcatattgtcctaaggcgttaccccaa**tca**gaggggtaccgagctcgaattcgtaatcatggtcatagctgtttcctgtgtgaaattgttatccgctcacaattccacacaacatacgagccggaagcataaagtgtaaagcctggggtgcctaatgagtga |

**Primers**

| **Name** | **Sequence** |
| --- | --- |
| **FW library** | aataggcgtatcacgaggc |
| **RV library** | agcgagtcagtgagcgag |

**Sequences of ssDNA and ssRNA (Figure 4)**

 Target sequences shown in blue

| **Sample** | **Sequence** |
| --- | --- |
| **ssDNA** | ggatcctaatacgactcactataggctgtccgatcgtataa**caggattccgcaatggggtt**accgcttaagcattaggggagctc |
| **ssRNA** | gcuguccgaucguauaa**caggauuccgcaaugggguu**accgcuuaagcauuaggggagcuc |

**sgRNA template (Figure 4)**

Target sequences shown in blue

| **sgRNA** | **Sequence** |
| --- | --- |
| sgRNA 20 nt S*treptococcus pyogenes*  ([WP_032464890.1](https://www.ncbi.nlm.nih.gov/protein/WP_032464890.1)) | taatacgactcactatagtcctaaggcgttaccccaagttttagagctagaaatagcaagttaaaataaggctagtccgttatcaacttgaaaaagtggcaccgagtcggtgctttt |
| sgRNA 18 nt S*treptococcus pyogenes*  ([WP_032464890.1](https://www.ncbi.nlm.nih.gov/protein/WP_032464890.1)) | taatacgactcactatagctaaggcgttaccccaagttttagagctagaaatagcaagttaaaataaggctagtccgttatcaacttgaaaaagtggcaccgagtcggtgctttt |
| sgRNA *Enterococcus faecium*  ([WP_119364770.1](https://www.ncbi.nlm.nih.gov/protein/1480008395)) | taatacgactcactataggtcctaaggcgttaccccaagttttagagctatgctgagaaatcaatatagcaagttaaaataaggctttgtccgtcatcagcttttttaaagcagcgctgttctcggcgcttttttt |
| sgRNA *Streptococcus thermophilus* (LMG 18311)  ([WP_011225725.1](https://www.ncbi.nlm.nih.gov/protein/499544942)) | taatacgactcactataggtcctaaggcgttaccccaagtttttgtactctcagaaatgcagaagctacaaagataaggcttcatgccgaaatcaacaccctgtcattttatggcagggtgtttt |
| sgRNA *Clostridium perfringens*  ([WP_003473526.1](https://www.ncbi.nlm.nih.gov/protein/489569047)) | taatacgactcactataggtcctaaggcgttaccccaagttatagttcctagtgaaaactagttactataacaaggcattaagccgtaaagtatcccctatgttcatttgaacctaggggtatcttttcattt |
| sgRNA *Staphylococcus aureus*  ([AYD60528.1](https://www.ncbi.nlm.nih.gov/protein/AYD60528.1)) | taatacgactcactataggtcctaaggcgttaccccaagttttagtactctggaaacagaatctactaaaacaaggcaaaatgccgtgtttatctcgtcaacttgttggcgagattt |
| sgRNA *Finegoldia magna*  ([WP_012290141.1](https://www.ncbi.nlm.nih.gov/protein/WP_012290141.1/)) | taatacgactcactataggtcctaaggcgttaccccaagtttgagaatgatgtaatgaaaattacatcatgagttcaaataaaagtttactcaaatcgcccgaaagagcccacattggtggactaaacaaatcttcggatttgttttttt |

**Customized Primers for NGS (Figure 5 and Supp Figure 8)**

Gene-specific sequences are indicated in blue

|  | **sgRNA (5'-3') (PAM)** | **Primer sgRNA (5'-3')** | **Primer sequence (5'-3')** |
| --- | --- | --- | --- |
| ***TYR* gene** | gcatcccgccagtcccaata (TGG) | FW: acaccgcatcccgccagtcccaatag | FW: ttcgatttgagtgccccaga |
|  |  | RV: aaaactattgggactggcgggatgcg | RV: ccttgatgggggctgcaat |
| ***OCA2* gene** | aggagctgccactgccatcg (GGG) | FW: acaccaggagctgccactgccatcgg | FW: aactctcggagtgagctgtg |
|  |  | RV: aaaaccgatggcagtggcagctcctg | RV: tctccagtgagagggaacagg |

|  | **Sequences** |
| --- | --- |
| ***TYR* gene** | TYREX1CRISP-F 5’-tcgtcggcagcgtcagatgtgtataagagacagtcaatggatgcactgcttgg-3’  TYREX1CRISP-R 5’- gtctcgtgggctcggagatgtgtataagagacagtcaatggatgcactgcttgg-3’ |
| ***OCA2* gene** | OCA2EX14CRISP-F 5’-tcgtcggcagcgtcagatgtgtataagagacagcgcctcccttatacgagcaa-3’  OCA2EX14CRISP-R 5’- gtctcgtgggctcggagatgtgtataagagacagttactgtgaagaggtggcgt-3’ |

|  | **Sequences** |
| --- | --- |
| ***TYR* gene** | TYREX1CRISP-F 5’-tcgtcggcagcgtcagatgtgtataagagacagtcaatggatgcactgcttgg-3’  TYREX1CRISP-R 5’- gtctcgtgggctcggagatgtgtataagagacagtcaatggatgcactgcttgg-3’ |
| ***OCA2* gene** | OCA2EX14CRISP-F 5’-tcgtcggcagcgtcagatgtgtataagagacagcgcctcccttatacgagcaa-3’  OCA2EX14CRISP-R 5’- gtctcgtgggctcggagatgtgtataagagacagttactgtgaagaggtggcgt-3’ |

**Traffic Light reporter assay (Supp Figure 9)**

|  | **Sequence** |
| --- | --- |
| **gRNA 1** (PAM GGG) | tacgcaaataagagctcacc |
| **gRNA 2** (PAM AGG) | aggtgagctcttatttgcgt |
| **SpCas9 U6 driven expression cassette** | ttaccgtaacttgaaagtatttcgatttcttggctttatatatcttgtggaaaggacgaaacaccgaggtgagctcttatttgcgtgttttagagctagaaatagcaagttaaaataaggctagtccgttatcaacttgaaaaagtggcaccgagtcggtgcttttttt |
